# Evolutionarily conserved coupling of lysosomal response sensing and disease-associated microglial reprogramming scales with pathological timescales

**DOI:** 10.64898/2026.11.14.766631

**Authors:** Yunlong Liang, Hua Yuan, Yayun Zhao, Li Guo, Juan wang, Liyi Chi, Yanhai Zhang

## Abstract

**Background:** Microglia transition into disease-associated microglia (DAM) during central nervous system (CNS) injury and neurodegeneration, yet the upstream intracellular logic driving this state conversion in vivo remains unclear. The lysosome acts as a signaling hub, but whether physiological phagocytic load universally initiates and quantitatively couples to DAM reprogramming across different pathological timescales is unknown.

**Methods:** We established a rigorous, unit-corrected analytical framework to overcome pseudo-replication in public single-cell data, anchoring statistical inferences at the level of independent biological units (animals, sequencing plates, or donors). We integrated six transcriptomic datasets spanning three murine injury/amyloidosis models and three large-scale human neurodegenerative cohorts (>320,000 microglial nuclei), alongside human Alzheimer’s disease (AD) GWAS summary statistics.

**Results:** We demonstrate that the MiT/TFE-regulated lysosomal biogenesis program temporally precedes DAM reprogramming and maintains an evolutionarily conserved baseline coupling even in uninjured homeostatic tissue. In human microglia, *MITF* and *TFE3* transcripts closely track DAM activation, whereas post-translationally regulated *TFEB* mRNA does not. Genetic enrichment analysis (MAGMA) reveals that common AD risk does not act upon the upstream lysosomal program, but is strongly anchored downstream at *APOE*. Furthermore, *Trem2*-knockout models demonstrate that disease-induced coupling amplification retains roughly half its magnitude even when the mature DAM subpopulation collapses. Cross-system comparisons reveal a state-transition synchrony model: disease-driven amplification of lysosomal–DAM coupling(Δ) scales with pathological timescales, showing strong amplification in acute murine injury and aggressive amyloidosis (5xFAD Δ = +0.252), but manifesting primarily as steady baseline maintenance across decades-long asynchronous human AD progression(SEA-AD Δ= +0.072).

**Conclusions:** Our study establishes the lysosomal stress–DAM coupling as an evolutionarily conserved pacemaking module for microglial state transitions, providing a quantitative framework for therapeutically targeting neuroinflammation and tissue repair.

## Background

As the resident myeloid immune cells of the central nervous system (CNS), microglia transition from homeostatic states into DAM—characterized by the upregulation of *Apoe*, *Trem2*, *Lpl*, and *Cst7*—in response to acute neural injury and chronic neurodegeneration [1–3]。This reprogramming process is essential for clearing myelin debris and amyloid deposits and for promoting tissue repair [4,5]. Although single-cell transcriptomics has extensively charted the molecular landscape of DAM, existing mechanistic studies have predominantly focused on downstream state amplifiers, such as TREM2 receptor signaling, lipid metabolic remodeling, and epigenetic modifications [1,2,4,5]. Within complex in vivo pathological microenvironments, however, the identity of the intracellular signals that act as upstream pacemakers to translate extracellular phagocytic burden into DAM transcriptional reprogramming remains elusive; specifically, quantitative in vivo evidence is lacking for an upstream intracellular program that precedes and tightly couples with the homeostatic-to-DAM transition across both injury models and chronic human diseases, even though the timing and efficiency of this transition directly dictate the trajectory of neural repair.

The lysosome serves not only as the terminal organelle for degrading phagocytic substrates, but also as a signaling hub that senses intracellular metabolic and mechanical states [6–8]. Following the engulfment of myelin debris or protein aggregates, alterations in luminal pH, calcium concentration, and membrane tension are sensed via v-ATPase, TRPML1 (encoded by *Mcoln1*), and the mechanosensitive channel TMEM63A, which promote the dephosphorylation and nuclear translocation of *MiT*/*TFE* family transcription factors (*TFEB*, *TFE3*, and *MITF)* to activate the downstream CLEAR lysosomal gene network [6–9]. Recent work has shown that elevated lysosomal membrane tension directly activates *TFEB* in peripheral macrophages [9]. Moreover, in primary lysosomal defect models—including mucopolysaccharidosis IIIA (MPS-IIIA) and progranulin (*GRN*)-deficient frontotemporal dementia—intrinsic lysosomal dysfunction is sufficient to drive DAM-like phenotypes via *MITF*/*TFE3*, while exogenous *Tfeb* overexpression ameliorates pathology in 5xFAD mice [10,11,12]. However, because these lines of evidence stem primarily from intrinsic genetic lysosomal defects or artificial overexpression, it remains unresolved whether lysosomal stress triggered by physiological injury-induced phagocytosis universally initiates and quantitatively couples to DAM conversion in vivo, and how this coupling evolves across acute insults versus decades-long chronic human neurodegeneration.

Here, we systematically tested whether the lysosomal stress program and the DAM program form a quantifiable coupling in vivo and how this coupling evolves across pathological timescales. To overcome cell-level pseudo-replication and batch-correction distortions of gene expression common to public single-cell reanalyses, we established a rigorous statistical unit-correction and batch-decoupled analytical framework (Fig. 1). We computed a 10-gene lysosomal program score (LYS10, capturing net *MiT*/*TFE* transcriptional output) and core DAM scores directly on unintegrated normalized expression matrices, anchoring formal statistical inferences at the level of independent experimental animals, sequencing plates, or human donors. By integrating murine peripheral nerve axotomy [13], central chemical demyelination [2], chronic amyloidosis (5xFAD and *Trem2* knockout) ^1^ three large-scale human neurodegenerative cohorts (Lee 2026 [14], Rexach 2024 [15], and SEA-AD [16]), and human AD genome-wide association study (GWAS) summary statistics, we delineate the temporal chronology, cross-species conservation, genetic anchoring, and timescale-scaling dynamics of lysosomal–DAM coupling. Together, these findings support our working model wherein injury-induced phagocytic load engages MiT/TFE factors (*TFEB*/*TFE3*/*MITF*) alongside lysosomal stress sensing, exhibiting temporal precedence and tight quantitative coupling with the establishment of an *Apoe*^+^ DAM repair state (Fig. 1b).

**Fig. 1.**
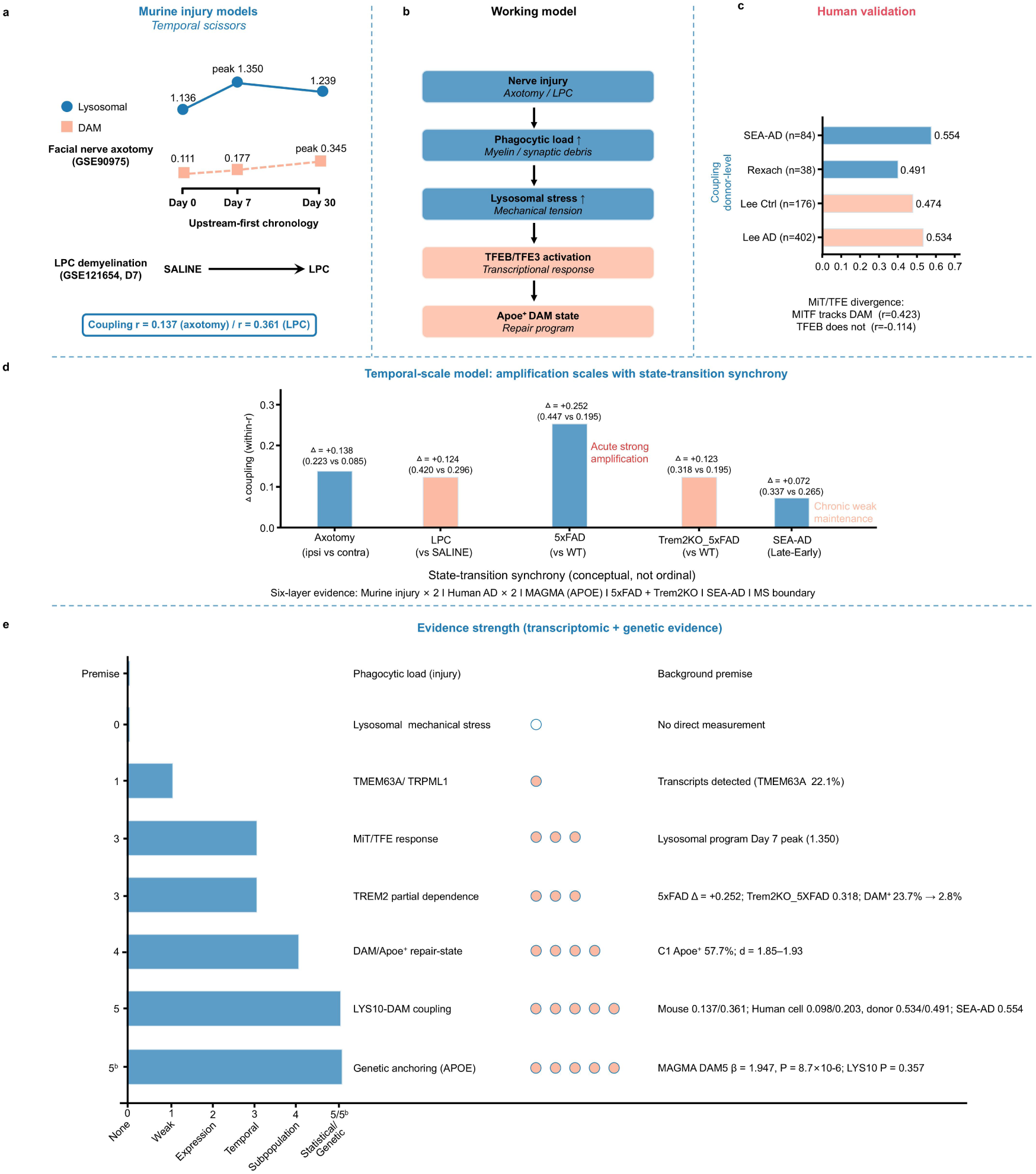
Overview of the study framework, working model, and evidence hierarchy. a, Temporal dynamics (temporal scissors) and pooled coupling strengths between the lysosomal program and the DAM program across two murine neural injury models (facial nerve axotomy, GSE90975; corpus callosum LPC demyelination, GSE121654). **b,** Schematic of the working model: neural injury-induced phagocytic load activates MiT/TFE transcription factors (*TFEB*/*TFE3*/*MITF*) via lysosomal stress, driving reprogramming into an *Apoe*^+^ DAM repair state. **c,** Donor-level coupling strengths across three human neurodegenerative cohorts (SEA-AD, Rexach 2024, and Lee 2026) and transcript-level regulatory divergence within the MiT/TFE family (*MITF* versus *TFEB*) relative to the DAM state. **d,** Cross-system comparison of coupling change magnitudes (Δ) across experimental and clinical systems, illustrating the “state-transition synchrony” timescale model (strong acute amplification in murine injury and 5xFAD versus weak chronic maintenance in human AD). **e,** Transcriptomic and genetic evidence hierarchy across components of the working model (levels 0–5 and the independent genetic tier 5b).

## Methods

### Data sources and cohort structure

Murine facial nerve axotomy dataset (GSE90975, discovery cohort) [13] : Generated on the mCEL-Seq2 platform, comprising 16 samples isolated from the ventral pontine facial nucleus of 12-week-old CX3CR1*^GFP/wt^*mice (*n* = 10 independent animals in total: 4 unoperated baseline mice at day 0; 3 operated mice each at days 7 and 30 post-axotomy, with each operated mouse contributing paired injured ipsilateral and uninjured contralateral samples; Supplementary Table 1). Samples were distributed across 4 sequencing plates (P1–P4), forming 14 batch-by-timepoint-by-laterality strata.

Murine corpus callosum demyelination dataset (GSE121654, external validation cohort) ^2^ : Generated on the Illumina NextSeq 500 platform, comprising FACS-purified CD11b*^+^* microglia harvested 7 days after stereotaxic corpus callosum injection of lysophosphatidylcholine (LPC, *n* = 3 independent P100 female mice) or saline (*n* = 3 independent P100 female mice; Supplementary Table 1).

Murine chronic amyloidosis dataset (GSE98969) [1]: Comprising brain microglia from 5xFAD transgenic mice (31 sequencing plates, 3,968 cells), littermate C57BL/6 wild-type controls (28 plates, 4,869 cells), *Trem2*-KO×5xFAD mice (6 plates), and *Trem2*-KO mice (3 plates), retaining plates with ≥20 cells (Supplementary Table 2). Because metadata lacked individual animal and litter identifiers, statistical inferences treated sequencing plates as the finest available independent units, supplemented by age-matched (6-month-old) sensitivity analyses.

Human neurodegenerative single-nucleus cohorts: Standardized .h5ad matrices were obtained from CZ CELLxGENE and the Allen Institute for Brain Science (with a pre-specified inclusion threshold of ≥ 20 nuclei per donor). (i) Lee 2026 cohort [14]: Encompassing 578 independent donors across the MSSM (*n* = 481) and RADC (*n* = 97) clinical centers (402 AD and 176 controls; 100,379 microglial nuclei; Supplementary Table 3). (ii) Rexach 2024 cohort [15] : Encompassing 38 qualifying independent donors sampled across the insula, primary motor cortex (BA4), and primary visual cortex (V1) (10 AD, 11 PSP, 8 Pick’s disease, and 9 controls; 82 donor-by-region units, 21,572 nuclei; Supplementary Table 3). (iii) SEA-AD cohort [16] : Comprising 236,002 microglial nuclei from 84 independent donors (median 2,460 nuclei per donor; all donors > 400 nuclei), spanning Braak stages 0 (*n* = 2), II (*n* = 4), III (*n* = 6), IV (*n* = 23), V (*n* = 34), and VI (*n* = 15), with a median age at death of 90 years (Supplementary Table 4). (iv) Human multiple sclerosis (MS) exploratory cohort (GSE118257) [17]: Comprising white matter lesion and control white matter myeloid nuclei across 5 donors (MS121, MS122, MS176, MS242, and SD48/16).

### Quality control, preprocessing, and batch-decoupling strategy

In murine datasets, ERCC spike-in controls and genes expressed in < 3 cells were removed, and high-quality cells with < 10% mitochondrial transcript fraction and sufficient detected genes (≥ 200 in GSE90975; ≥ 500 in GSE121654) were retained (1,437 cells × 13,086 genes in GSE90975; 5,325 cells × 18,735 genes in GSE121654). Following log1*p* normalization, the top 2,000 highly variable genes were selected for principal component analysis (PCA). Core batch-decoupling principle: Harmony integration (v0.0.9) was applied exclusively to the 30-dimensional PCA embedding (X_pca_harmony) for Leiden clustering and UMAP visualization; all gene module scores, differential expression analyses, and regression models were computed directly on the unintegrated, normalized expression matrix (adata.X), strictly preventing batch-integration algorithms from artificially introducing inter-gene covariation. Human cohorts used officially released standardized expression matrices directly, extracting per-nucleus total UMI counts from the raw count layer (raw.X) as sequencing depth covariates.

### Transcriptomic functional module scoring

Module scores were computed using the score_genes function in Scanpy (v1.12.3), which partitions all genes into n_bins = 25 expression bins by mean abundance and subtracts the mean expression of ctrl_size = 50 randomly sampled control genes per bin:

Lysosomal program score (LYS10): Composed of 10 core lysosomal biogenesis and hydrolytic enzyme genes (mouse: *Lamp1*, *Ctsb*, *Ctsd*, *Ctsl*, *Ctss*, *Atp6v0d1*, *Sqstm1*, *Npc1, Gba,* and *Psap*; human orthologous genes, where *GBA* maps to annotated *GBA1*). To prevent analytical circularity, upstream echanosensitive/calcium channels (*Mcoln1* and *Tmem63a*), low-detection transcription factors themselves (*Tfeb* and *Tfe3*), and the downstream effector *Apoe* were strictly excluded from LYS10.

Core DAM scores (DAM5 and DAM4): DAM5 comprised *Apoe*, *Trem2*, *Lpl*, *Cst7,* and *Ccl5* (and human orthologs); to test *Apoe*-independence, we constructed a 4-gene version excluding *Apoe*/*APOE* (DAM4), supplemented by distribution-free rank-percentile scoring for sensitivity validation.

Injury-repair signature (REPAIR12) and C1 marker projection: The 12-gene repair signature comprised *Apoe*, *Lgals3bp, H2-K1*, *Ifitm3*, *H2-D1*, *Ifi27l2a*, *Rps21*, *Rpl36a*, *Ccl5, Cplx2*, *Gabra3,* and *Fcgr2b* (mapped to human orthologs as REPAIR12, alongside *Apoe*-removed [no*Apoe*] and *Apoe*/*Ccl5*-removed [noAC] variants). The top 15 C1 subpopulation markers (Supplementary Table 5) and their *Apoe*/*Cst7*-removed de-overlapped variants were projected onto external datasets, evaluating shared-gene attenuation via the retention ratio (*r_de−overlapped_/r_full_*). For the memory-intensive SEA-AD matrix, we implemented a sparse CSR chunked scanner that reproduced standard in-memory score_genes outputs at *r* = 0.984–0.990 on a validation subset.

### Statistical modeling and independent unit-correction hierarchy

To overcome pseudo-replication inflation in single-cell reanalyses, we implemented a four-tiered complementary statistical inference framework.

### Cluster-robust linear regression (Cluster-robust OLS, primary test)

In murine injury models, we fitted the interaction model:

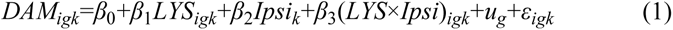

Using the Liang–Zeger sandwich variance estimator clustered across the *G* = 14 strata in GSE90975 or *G* = 6 independent animals in GSE121654 with the finite-sample correction factor *c*, two-sided *P*-values for the interaction term *β_3_* were evaluated against a conservative *t*(*G* - 1) distribution; for GSE121654, we additionally performed an exhaustive wild cluster bootstrap across all 2^6^ = 64 sign permutations (see Supplementary Methods for details).

### Linear mixed-effects models (Corroborative test)

Treating sequencing plates, individual animals, or human donors (e.g., the 38 donors in the Rexach cohort) as random intercepts.

### Within-unit correlation and Fisher *z*-pooling

Single-cell Pearson correlation coefficients *r_k_* were computed within each independent stratum, sequencing plate, or human donor, transformed via Fisher’s *z* = *artanh(r_k_)*, and pooled across units using inverse-variance weights (*w_k_=n_k_*−3); alternatively, plate or donor means were compared directly as independent observations using two-sided Welch’s *t*-tests, Mann–Whitney *U*-tests, and one-sample *t*-tests.

### Leave-one-unit-out cross-validation

Units (sequencing plates or individual animals) were removed one at a time, and the estimate from the worst-performing fold is reported.

### Human SEA-AD cohort

Within-donor correlation coefficients were converted to Fisher z scores (within-z) and fitted by WLS regression against continuous Braak stage, age, sex and log10(nucleus number), weighted by nucleus number. Pairwise comparisons across coarse stage groups (early, middle and late) were performed using two-sided Mann–Whitney *U* tests with Bonferroni correction for three comparisons.

### Pathway enrichment and MAGMA genetic analyses

Two-sided Fisher’s exact tests were performed with Enrichr/GSEApy (v1.3.1) based on locally archived snapshots of the KEGG_2019_Mouse and GO_Biological_Process_2023 gene set collections; false discovery rate was controlled by the Benjamini–Hochberg method (adjusted *P* < 0.05). Gene set enrichment analysis was performed with MAGMA (v1.10) using AD GWAS summary statistics from Bellenguez et al. (2022) (GCST90027158, excluding 23andMe). SNPs were mapped to GRCh37/NCBI37.3 genomic coordinates by rsID (18,349 genes mapped in total), with the 1000 Genomes Project European population used as the linkage disequilibrium reference panel. Six gene sets (LYS10, DAM5, REPAIR8, GO_LYSOSOME, GO_AUTOPHAGY and the positive control KEGG_AD) were tested jointly in a conditional model, with Bonferroni significance threshold set at *P* < 0.05/6 = 0.0083; a LOO sensitivity analysis was additionally performed for DAM5.

### Software and reproducibility

All analyses were performed in Python (v3.10) using Scanpy (v1.12.3), harmonypy (v0.0.9), statsmodels (v0.14.6), SciPy (v1.18.0), GSEApy (v1.3.1) and MAGMA (v1.10). All analysis scripts, configuration files and intermediate results have been publicly archived on Zenodo (10.5281/zenodo.23259475) to Unless otherwise stated, all statistical tests were two-sided, with significance level set at α = 0.05.

## Results

### Neural injury induces an *Apoe*^+^ repair-state microglial subpopulation (C1) coupling phagocytic degradation with innate immunity

To capture the terminal effector state of the DAM transition during acute neural injury, we first analyzed the murine facial nerve axotomy dataset(GSE90975, *n* = 10 independent mice across 4 sequencing plates, retaining 1,437 quality-controlled microglia**) [**13]. To prevent batch correction from distorting gene expression values, the Harmony algorithm was applied exclusively to the dimensionality-reduced embedding for unsupervised Leiden clustering and visualization, while all gene module scores were computed independently on the unintegrated expression matrix. Following batch correction, cells from the four sequencing plates mixed uniformly (Fig. 2a, b), and cluster boundaries as well as coupling estimates remained highly stable across Harmony penalty parameters *θ*∈{0.5,2.0,5.0} (Supplementary Fig. 1).

**Fig. 2.**
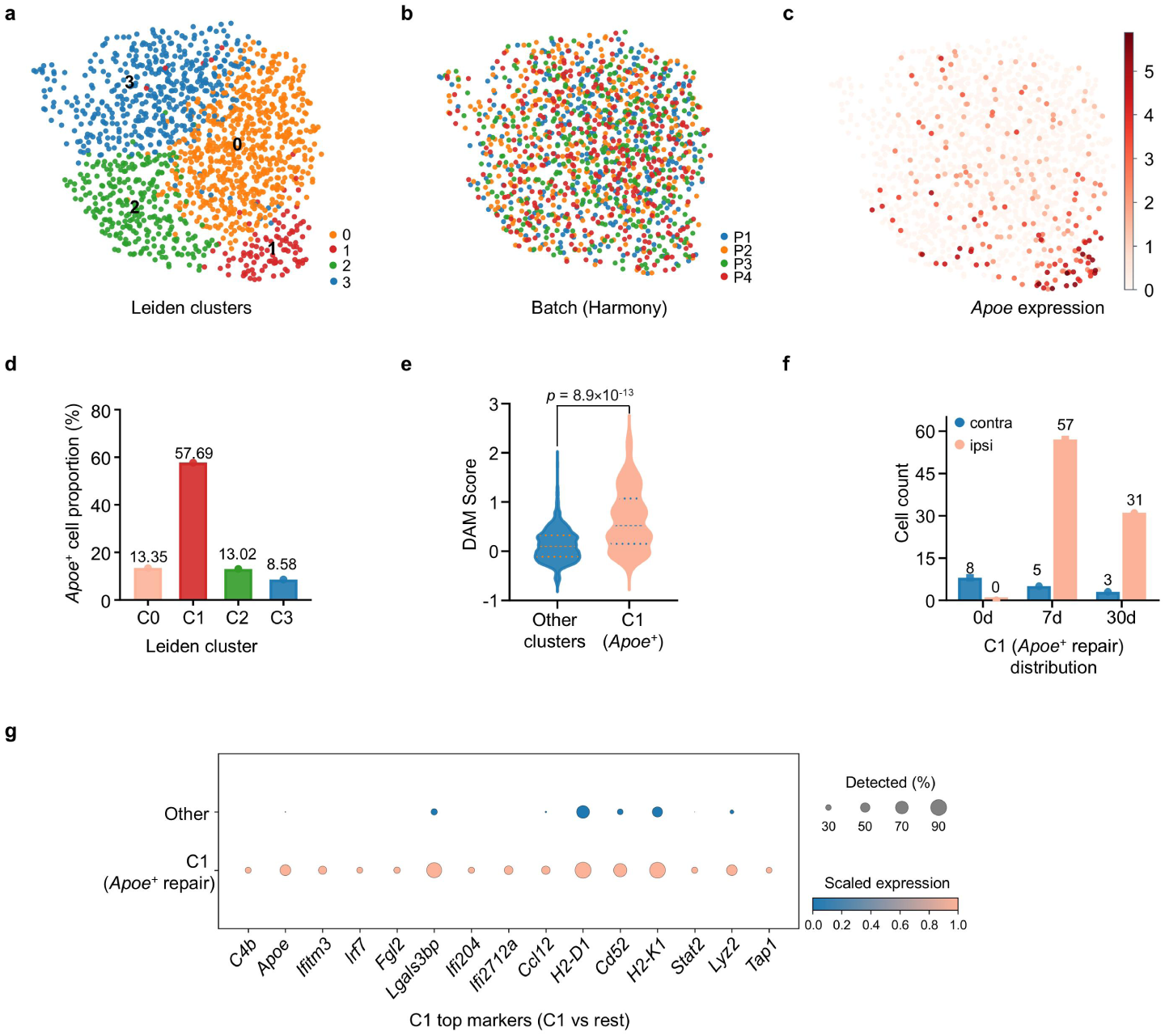
Identification and transcriptomic characterization of the repair-state microglial subpopulation C1 in the murine facial nerve axotomy model (GSE90975, *n* = 10 independent mice, 1,437 cells). **a**, UMAP visualization of unsupervised Leiden clusters (clusters C0–C3). **b**, UMAP distribution after Harmony batch correction, colored by sequencing plate (P1–P4). **c**, Spatial distribution of *Apoe* expression levels on the UMAP embedding. **d,** Proportion of *Apoe*^+^ cells (%) across Leiden clusters. **e**, Violin plot comparing DAM scores between subpopulation C1 and the remaining clusters (two-sided *t*-test, *P* = 8.9 × 10^-13^). **f**, Cell count distribution of subpopulation C1 across time points (days 0, 7, and 30) and anatomical sides (contralateral, contra; injured ipsilateral, ipsi). **g**, Dot plot of representative marker gene expression in subpopulation C1 versus the remaining clusters (dot size indicates the percentage of cells with detected expression; color scale indicates standardized expression level).

Within the clustering manifold, we identified a distinct subpopulation (designated C1, comprising 104 cells) highly enriched for *Apoe* transcripts (Fig. 2c). Although Apoe expression was excluded from the clustering computation itself, the proportion of *Apoe*^+^ cells within C1 reached 57.69% (60/104), far exceeding the 8.58%–13.35% observed in the remaining clusters (Fig. 2d), and its mean DAM score (0.614) was significantly higher than that of the remaining clusters (0.128, two-sided *t*-test *P* = 8.9 ×10^-13^; Fig. 2e). Spatiotemporally, C1 exhibited a pronounced skew toward the injured side: 88 of the 104 cells originated from the injured ipsilateral side (57 at day 7 and 31 at day 30 post-axotomy), whereas C1 was completely absent on the ipsilateral side at the day 0 baseline (Fig. 2f). Scoring against published microglial reference signatures confirmed that C1 displayed significantly elevated DAM (*P* = 7.4 × 10^-17^) and interferon-responsive microglia (IRM; *P* = 1.6 ×10^-7^) scores alongside a significantly reduced homeostatic score (*P* = 1.8 × 10^-8^; Supplementary Fig. 2).

Differential expression analysis (Fig. 2g and Supplementary Table 5) revealed that, beyond marked Apoe upregulation (log2FC = 4.36), C1 coordinately activated three functional modules: (1) a lysosomal cathepsin axis (*Ctsl*, *Ctsb*, and *Ctss*; log_2_FC = 1.10–1.78); (2) an MHC class I antigen presentation axis (*H2-D1*, *H2-K1*, and *B2m*); and (3) an interferon and innate immune response module (*Ifitm3*, *Irf7*, *Fgl2*, and *C4b*). Pathway enrichment and sensitivity analyses removing MHC-I genes (*H2-K1* and *H2-D1*) confirmed that this repair signature organizes around a phagolysosomal and type II interferon/innate immune core independent of antigen presentation genes or data-processing workflows (Supplementary Fig. 3 and Supplementary Table 6). Conditional independence analysis demonstrated that after controlling for C1 membership, the main effect of the lysosomal program on the DAM score remained highly significant (+0.084, *P* = 1.9 × 10^-6^); excluding all C1 cells reduced the overall pooled correlation *r* from 0.137 to 0.076 (retaining 55.5%, *P* = 0.006) but completely abolished the ipsilateral injury amplification interaction term (*P* = 0.810; Supplementary Table 7a,b). This indicates that baseline lysosomal–DAM coupling is broadly distributed across the microglial population, whereas subpopulation C1 represents the terminal effector state that harbors injury-induced lateral amplification.

### *Apoe^+^* cells expand robustly within individual batches, and lysosomal program activation temporally precedes DAM reprogramming

Across the entire facial nerve axotomy dataset, *Apoe*^+^ microglia (216 cells, 15.0%) exhibited a mean DAM score approximately tenfold higher than that of *Apoe*^-^ cells (0.689 versus 0.070, *P* = 1.1 × 10-51; Fig. 3a), and single-cell *Apoe* expression correlated strongly with the DAM score (Pearson *r* = 0.674, *P* = 3.8 × 10^-191^; Fig. 3b). The proportion of *Apoe*^+^ cells increased stepwise over time on the injured ipsilateral side—rising from 7.1% at day 0 to 24.4% at day 7 and 30.8% at day 30—while remaining at baseline levels of 6.8%–8.8% on the contralateral side (Fig. 3c). Pseudobulk comparisons treating individual animals as independent statistical units confirmed that the ipsilateral *Apoe*^+^ proportion was significantly higher than the contralateral proportion at both day 7(24.3% ± 1.3% versus 8.8% ± 5.7%, *P* = 0.041) and day 30 (30.6% ± 6.9% versus 6.7% ± 2.6%, *P* = 0.026), with a significant two-way regression interaction term (+10.78, *P* = 0.012). Crucially, within all five testable sequencing-plate-by-timepoint strata, ipsilateral elevations in both DAM scores and *Apoe*^+^ cell proportions were directionally consistent and individually reached statistical significance in the majority of strata (Fig. 3d, e), ruling out batch confounding. Cross-module comparisons showed that *Apoe*^+^ cells coordinately upregulated the *TFEB* CLEAR lysosomal, phagocytosis, and lipid metabolism modules (all *P* < 1 × 10^-4^), whereas the pro-inflammatory module showed only a marginal difference (*P* = 0.053; Fig. 3f, g).

**Fig. 3.**
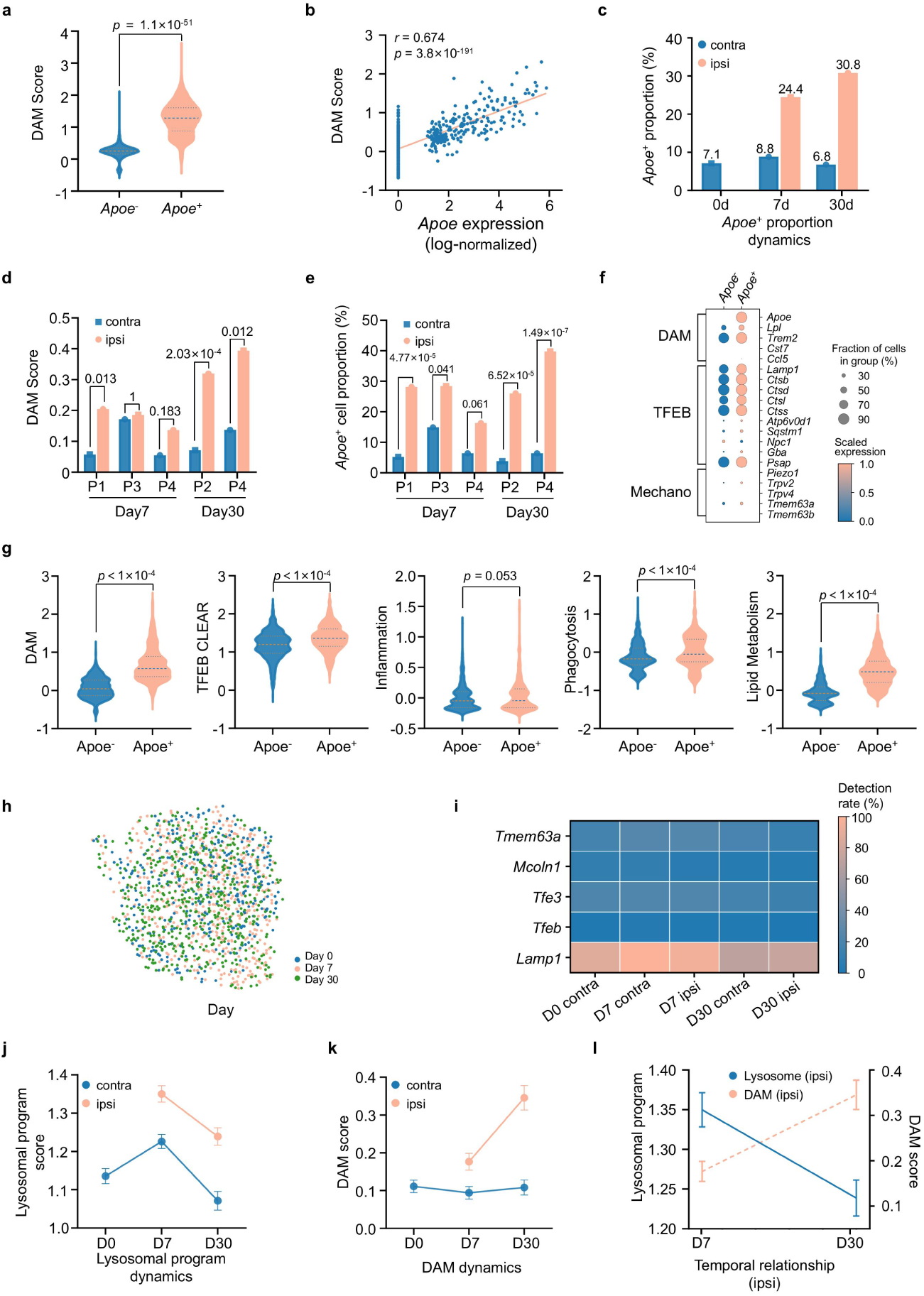
Functional localization, within-batch replication, and temporal dynamics of *Apoe*⁺ microglia (GSE90975, *n* = 10 independent mice, 1,437 cells, including 216 *Apoe*⁺ cells). **a**, Violin plot comparing DAM scores between *Apoe*^-^ and *Apoe*⁺ cells. **b**, Scatter plot and linear fit of single-cell log-normalized *Apoe* expression against DAM scores. **c**, Dynamics of *Apoe*⁺ cell proportions across time points (days 0, 7, and 30) on the contralateral (contra) and injured ipsilateral (ipsi) sides. **d, e**, Within-plate comparisons of contralateral versus ipsilateral DAM scores (d) and *Apoe*⁺ cell proportions (e) across individual sequencing plates (P1–P4) at days 7 and 30 post-axotomy (numbers above bars indicate *P*-values). **f**, Dot plot of representative DAM, *TFEB* lysosomal, and mechanosensing (Mechano) gene expression in *Apoe*^-^ and *Apoe*⁺ cells. **g**, Violin plots comparing module scores across five functional programs between *Apoe*^-^ and *Apoe*⁺ cells. **h**, UMAP distribution colored by sampling time (Day 0, Day 7, and Day 30). **i**, Heatmap of cellular detection rates for *TFEB*-axis and mechanosensing node genes across timepoint-by-laterality groups. **j, k**, Temporal trajectories of contralateral and ipsilateral lysosomal program scores (j) and DAM scores (k) (mean ± s.e.m.). **l**, Inverse crossover temporal trajectories of the ipsilateral lysosomal program score (blue solid line, left axis) and DAM score (orange dashed line, right axis) between days 7 and 30 post-axotomy.

Diffusion pseudotime analysis further supported a continuous transition trajectory from homeostatic to *Apoe*^+^ states (Fig. 3h and Supplementary Fig. 4). Temporal dynamics revealed a striking chronological “scissors” pattern between upstream and downstream programs (Fig. 3i, j–l). Following injury, cellular detection rates of lysosomal membrane tension- and calcium-sensing node genes (*Lamp1*, *Tmem63a*, and *Mcoln1*) and transcription factors (*Tfe3* and *Tfeb*) rose broadly by day 7 and remained elevated through day 30 (for example, *Lamp1* increased from a 74.7% baseline to 81.5%–85.2%, and Tmem63a rose from 20.5% to 25.9%–26.1%; Fig. 3i). At the functional module level, the ipsilateral lysosomal program score peaked rapidly at day 7 (1.350, versus 1.136 at baseline) before declining to 1.239 at day 30 (Fig. 3j); conversely, the ipsilateral DAM score showed only a modest elevation at day 7 (0.177, versus 0.111 at baseline) before ascending continuously to reach its peak at day 30 (0.345; Fig. 3k). Between days 7 and 30, the decline of the lysosomal program and the ascent of the DAM program formed a distinct inverse crossover trajectory (Fig. 3l). This population kinetic profile—characterized by an early upstream peak and decline followed by lagged downstream accumulation—provides strong temporal support for a model in which lysosomal stress sensing precedes and drives the establishment of the DAM repair state.

### Establishment of lysosomal–DAM coupling and statistical unit correction across murine acute neural injury models

To rigorously test quantitative coupling between the LYS10 lysosomal program—excluding *Apoe* and mechanosensitive channel genes to avoid circularity—and the DAM score, we performed cluster-robust ordinary least-squares (OLS) regression across the 14 batch-by-timepoint-by-laterality clusters in the facial nerve axotomy discovery cohort (GSE90975). This revealed a lysosomal program-by-ipsilateral interaction coefficient of +0.177 (conservative *t*(13)-based *P* = 0.0028, 95% CI: 0.061–0.292; Fig. 4a and Supplementary Table 8), identical in point estimate to naive OLS and demonstrating that correcting for pseudo-replication does not abolish the effect. Linear mixed-effects models treating sequencing plates or individual animals as random intercepts (coefficient +0.177, *P* = 0.0011) and leave-one-plate-out cross-validation (worst-fold coefficient +0.159, *P* = 0.012) confirmed the robustness of this interaction (Fig. 4b and Supplementary Fig. 5a, c–f). Stratified slope analysis showed that the overall regression slope on the injured ipsilateral side (+0.290, *P* = 8.2 × 10^-7^) was approximately 3.6-fold steeper than the contralateral baseline slope (+0.081, *P* = 1.9 × 10^-8^; Fig. 4c).

**Fig. 4.**
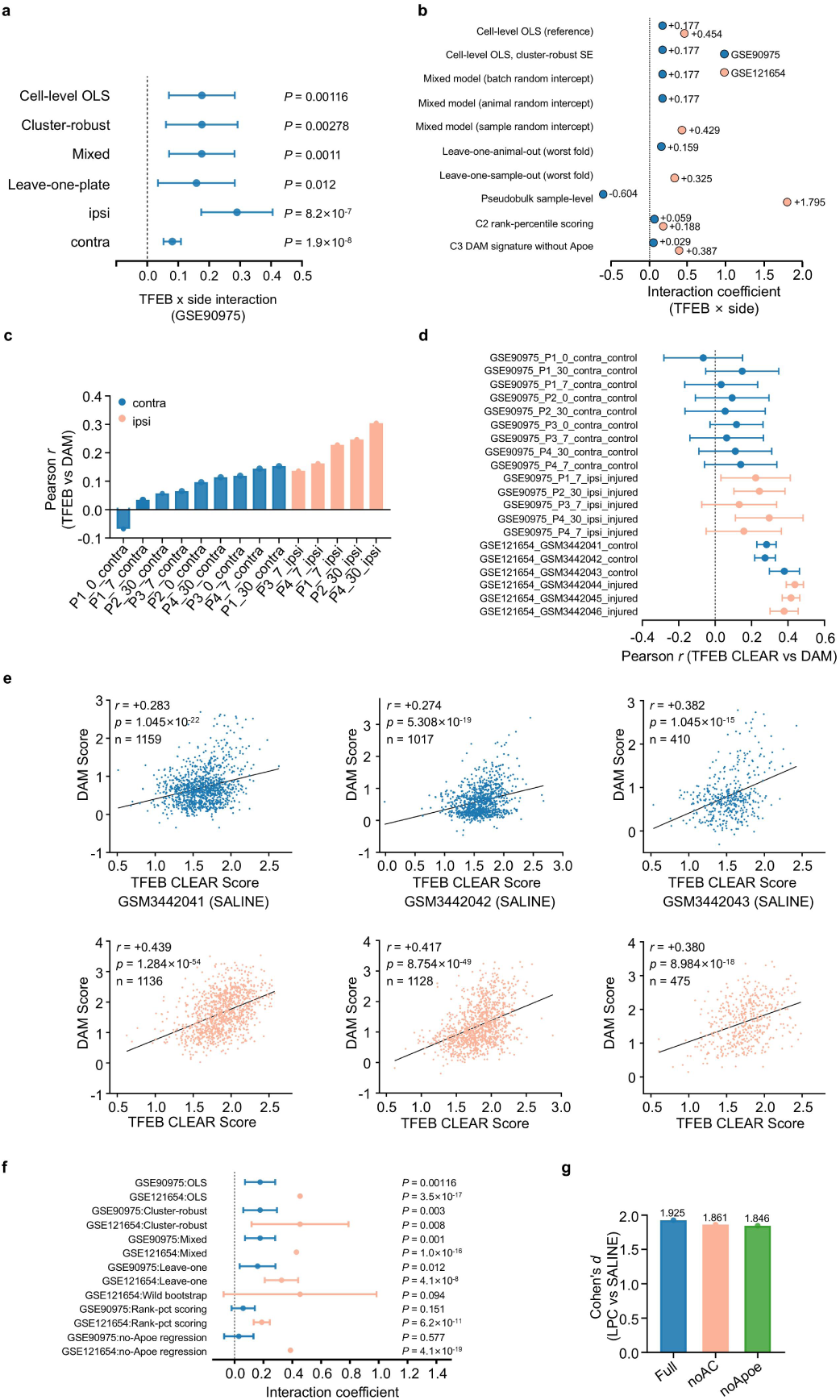
Validation of lysosomal–DAM coupling and statistical unit correction across two murine neural injury models (GSE90975, *n* = 10 independent mice across 14 strata; GSE121654, *n* = 6 independent mice). **a**, Forest plot of lysosomal program-by-laterality interaction coefficients across statistical model specifications and stratified ipsilateral (ipsi) and contralateral (contra) slopes in GSE90975 (error bars represent 95% CIs). **b**, Summary comparison of interaction point estimates across modeling and scoring specifications in both datasets. **c**, Bar plot of within-stratum coupling correlation coefficients (*r*) across the 14 contralateral and injured ipsilateral strata in GSE90975. **d**, Forest plot of within-group Pearson *r* values across all 20 batch/sample-by-condition groups in the two datasets (error bars represent 95% CIs). **e,** Single-cell scatter plots and linear fits of lysosomal program scores versus DAM scores across the six individual mice in GSE121654 (saline controls in blue, *n* = 3; LPC demyelination in orange, *n* = 3). **f**, Forest plot of interaction coefficients across all robustness specifications (including cluster-robust regression, mixed-effects models, leave-one-out cross-validation, wild cluster bootstrap, rank-percentile scoring, and *Apoe*-removed regression) in both datasets. **g**, Cell-level effect sizes (Cohen’s *d*) comparing LPC versus saline groups in GSE121654 across three repair signature versions: full signature (Full), *Apoe*- and Ccl5-removed (noAC), and *Apoe*-removed (no*Apoe*).

We next performed external validation in the independent corpus callosum focal LPC demyelination dataset (GSE121654; day 7 post-injection, *n* = 3 independent female mice per group in saline and LPC, totaling 5,325 FACS-purified microglia). Following Fisher *z*-transformation and pooling, coupling strength rose significantly from *r* = 0.296 (95% CI: 0.26–0.330) in the saline control group to *r* = 0.420 (95% CI: 0.389–0.450) in the LPC injury group, with an overall pooled correlation of *r* = 0.361across all samples (Fig. 4d, Supplementary Table 9, and Supplementary Fig. 6). Within all six individual mice, single-cell lysosomal–DAM correlations were highly significant (Pearson *r* = 0.274–0.439, all *P* < 10^-14^; Fig. 4e), and cluster-robust regression clustering by individual animal (interaction +0.454, conservative *t*(5) *P* = 0.008), animal random-intercept mixed-effects modeling (+0.429, *P* = 1.0 × 10^-16^), and leave-one-animal-out cross-validation (worst fold +0.325, *P* = 0.047) yielded highly concordant results (Fig. 4b, f, Supplementary Table 8, and Supplementary Fig. 7). Random-effects meta-analysis across the two murine injury models confirmed a pooled interaction coefficient of +0.301 (*P* = 0.029) and a Fisher *z*-pooled correlation of +0.251 (*P* = 0.032; Supplementary Table 10a, b and Supplementary Fig. 8).

To rule out potential collinearity arising from high *Apoe* expression within the DAM signature, we constructed an *Apoe*-removed DAM4 score. Upon removing *Apoe*, baseline coupling remained significant in both datasets (GSE90975 pooled *r* = +0.092, contralateral *P* = 0.034, ipsilateral *P* = 0.0037; GSE121654 pooled *r* = +0.293, LPC regression interaction +0.387, *P* = 4.1 × 10^-19^; Fig. 4b, f and Supplementary Table 9); association decomposition indicated that non-*Apoe* components carried 75.6% of the total coupling signal (Supplementary Table 11). Furthermore, projecting the C1 top markers or the 12-gene repair signature defined in the discovery cohort (alongside de-overlapped variants removing *Apoe*/*Ccl5*) onto the LPC demyelination and 5xFAD models achieved significant separation between injured/diseased and control groups at the animal or plate level (complete 3-versus-3 animal-level rank separation in LPC versus saline, Cohen’s *d* = 1.85–1.93, Welch’s *t*-test *P* = 0.008–0.012; Fig. 4g, Supplementary Figs. 8–10 and 14a–f, and Supplementary Table 12). Quintile dose–response analyses, three-level variance decomposition, sequencing depth (UMI) adjustments, negative-control outcome tests, replacement with external independent gene sets (GO_Lysosome and Hammond repair signatures), and E-value sensitivity analyses for unmeasured confounding further verified that this coupling follows a monotonic dose–response relationship (rising significantly from the fourth quintile onward, trend *P* = 3.3 × 10^-4^) and is not driven by Simpson’s paradox or technical artifacts (Supplementary Figs. 11–13 and Supplementary Table 10,11,13 and 14).

### Cross-species conservation of lysosomal–DAM coupling and MiT/TFE family regulatory divergence in human neurodegeneration

To test whether the coupling mechanism identified in mice translates to human disease, we analyzed three independent post-mortem human brain single-nucleus RNA-seq (snRNA-seq) cohorts: Lee 2026 (578 donors, 100,379 microglial nuclei) ^14^, Rexach 2024 (38 donors spanning AD, Pick’s disease, progressive supranuclear palsy [PSP], and controls, 21,572 nuclei) ^15^, and SEA-AD (84 donors, 236,002 nuclei) ^16^. UMAP projections visually illustrated the co-localization of percentile-scaled lysosomal and DAM scores (Fig. 5a); to prevent inflated precision from pseudo-replication across large nuclei counts, we anchored formal statistical inferences strictly at the independent donor level.

**Fig. 5.**
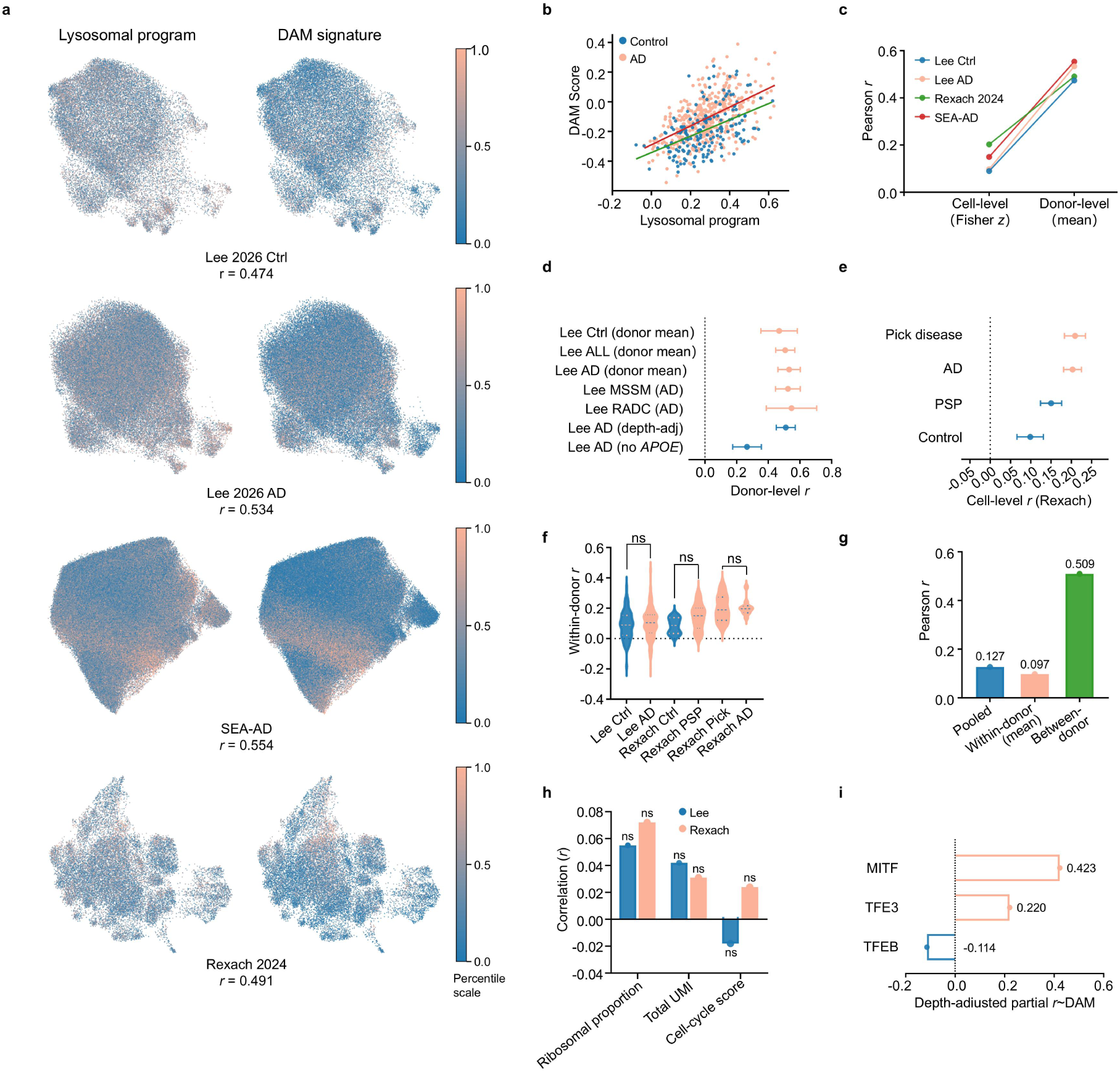
Cross-species validation of lysosomal–DAM coupling in human neurodegenerative cohorts (Lee 2026, *n* = 578 independent donors; Rexach 2024, *n* = 38 independent donors; SEA-AD, *n* = 84 independent donors). **a**, UMAP embeddings across four cohort subsets overlaid with percentile scores of the lysosomal program (left) and DAM signature (right), annotated with donor-level correlation coefficients (*r*). **b**, Scatter plot and linear fits of donor-mean lysosomal scores versus DAM scores in control (blue) and AD (red) donors from the Lee 2026 cohort. **c**, Systematic comparison of cell-level (Fisher *z*-pooled) versus donor-level (donor-mean or mixed-model) coupling correlation coefficients (*r*) across cohorts. **d**, Forest plot of donor-level correlation coefficients across diagnostic groups, clinical center subcohorts, depth-adjusted specifications, and *APOE*-removed signatures in the Lee cohort (error bars represent 95% CIs). **e**, Forest plot of cell-level pooled correlation coefficients across diagnostic groups in the Rexach 2024 cohort (error bars represent 95% CIs). **f**, Violin plots of within-donor correlation coefficients (within-donor *r*) across diagnostic groups in the Lee and Rexach cohorts. **g**, Three-level correlation decomposition in the Lee cohort. **h,** Technical negative-control associations in the Lee (blue) and Rexach (orange) cohorts (ns, not significant). **i,** Depth-adjusted donor-level partial correlation coefficients between MiT/TFE family transcripts and the DAM score in the Lee cohort.

Across all three human cohorts, both cell-level (donor-stratified Fisher *z*-pooling) and donor-level (donor-mean correlation or random-intercept mixed-effects modeling) analyses confirmed robust positive coupling (Fig. 5a–d and Supplementary Table 15 and 16). At the cellular level, the overall pooled correlation was *r* = 0.096 (in the Lee cohort (0.098 in AD and 0.090 in controls) and *r* = 0.174 in the Rexach cohort, remaining positive across all four diagnostic groups along a gradient from Pick’s disease (0.209) and AD (0.203) to PSP (0.150) and controls (0.099; Fig. 5e), mirroring the severity of cortical tau pathology across these tauopathies. Notably, donor-level coupling was systematically stronger than cell-level coupling: *r* = 0.474 in Lee controls, *r* = 0.534in Lee AD (0.527 and 0.560 in the MSSM and RADC clinical center subcohorts, respectively; Fig. 5c, d), *r* = 0.554 in SEA-AD, and a standardized mixed-model coefficient of +0.491 in Rexach (*P* = 6.2 × 10^-12^P = 6.2 × 10^-12^; Fig. 5c). Three-level correlation decomposition revealed that between-donor covariation (Lee cohort *r* = 0.509) substantially exceeded mean within-donor cellular covariation (mean *r* = 0.097; Fig. 5f, g), indicating that individuals with higher baseline lysosomal tone concurrently mount stronger overall DAM activation. Adjusting for sequencing depth or brain region covariates fully preserved donor-level partial correlations (retaining 101%–107.5%), removing *APOE* retained approximately half of the donor-level coupling (Lee AD *r* = 0.267; Fig. 5d), and three technical negative-control outcomes (ribosomal gene fraction, total UMI count, and cell-cycle score) yielded near-zero effect sizes (Fig. 5h and Supplementary Table 16 and 17).

These large-scale human cohorts also uncovered a striking transcript-level functional divergence within the MiT/TFE transcription factor family (Fig. 5i, Supplementary Table 18, and Supplementary Fig. 15).

At the donor level in the Lee cohort after adjusting for sequencing depth, *MITF* transcript abundance correlated strongly with the DAM score (partial *r* = 0.423, *P* = 1.7 × 10^-26^) and *TFE3* showed a moderate positive correlation (*r* = 0.220, *P* = 9.6 × 10^-8^), whereas *TFEB* mRNA abundance failed to track the DAM state and instead exhibited a weak inverse correlation (*r* = −0.114, *P* = 0.0063; a modest unadjusted positive association with the lysosomal score, *r* = +0.104, was driven entirely by sequencing depth confounding and reversed to *r* = −0.163) after depth adjustment). This finding aligns with classic biochemical mechanisms: *TFEB* activation is governed primarily post-translationally via mTORC1- and calcineurin-mediated dephosphorylation and nuclear translocation rather than steady-state mRNA synthesis, whereas *MITF* is transcriptionally upregulated in human neurodegenerative microglia [6–8,10,11,14,18]. Consequently, our 10-gene CLEAR lysosomal score (LYS10) accurately captures the shared downstream transcriptional de-repression of the *MiT*/*TFE* family, overcoming the blind spot of monitoring *TFEB* mRNA alone. Regarding disease modulation of the coupling slope, AD status and advancing age in the Lee cohort primarily shifted microglia upward along the coupling axis (both main effects *P* < 0.001) without altering the slope itself (interaction *P* = 0.476; Fig. 5b), whereas the smaller Rexach cohort exhibited a significant disease interaction (*b* = +0.432, *P* = 0.008; Supplementary Table 16), suggesting that slope modulation in chronic human disease may be sensitive to cohort heterogeneity and regional sampling.

### Common AD genetic risk is anchored downstream at *APOE*, and coupling amplification in the 5xFAD model is partially independent of *TREM2*

To dissect the genetic architecture of the lysosomal–DAM coupling axis, we performed conditional gene-set enrichment analysis using MAGMA (v1.10) with summary statistics from the large-scale 2022 Bellenguez et al. AD GWAS (up to 85,934 cases and 401,577 controls; Fig. 6a–c and Supplementary Table 19). Genome-wide significant loci accurately anchored to the *APOE* gene cluster (*Z* = 7.53) alongside *BIN1*(*Z* = 9.68), *ABCA7* (*Z* = 9.34), *SORL1* (*Z* = 8.01), and the positive-control KEGG Alzheimer’s disease pathway was significantly enriched (*β* = 0.209, *P* = 0.0032; Fig. 6a). In a joint conditional model, the downstream DAM5 signature exhibited strong genetic risk enrichment (*β* = 1.947, *P* = 8.7 × 10^-6^, surpassing the Bonferroni threshold of *α* = 0.0083), whereas the upstream lysosomal program (LYS10: *β* = 0.106, *P* = 0.357) and reference autophagy/lysosome sets (GO_AUTOPHAGY *P* = 0.372; GO_LYSOSOME *P* = 0.083) showed no enrichment. Leave-one-gene-out (LOO) analyses revealed that DAM5 genetic enrichment was driven almost entirely by *APOE*: removing *APOE* abolished enrichment (*β* = 0.389, *P* = 0.257), whereas removing any of the other four genes (*TREM2*, *LPL*, *CST7*, or *CCL5*) preserved significance(*P* = 0.004–0.034; Fig. 6b, c). Notably, the 8-gene repair signature (REPAIR8)—which also contains *APOE*—failed to reach significance (*P* = 0.071)), confirming that *APOE* inclusion alone does not artificially manufacture set-level significance. Thus, common late-onset AD genetic risk variants do not directly modify the upstream lysosomal sensor, but instead anchor with high specificity at the downstream effector *APOE*.

**Fig. 6.**
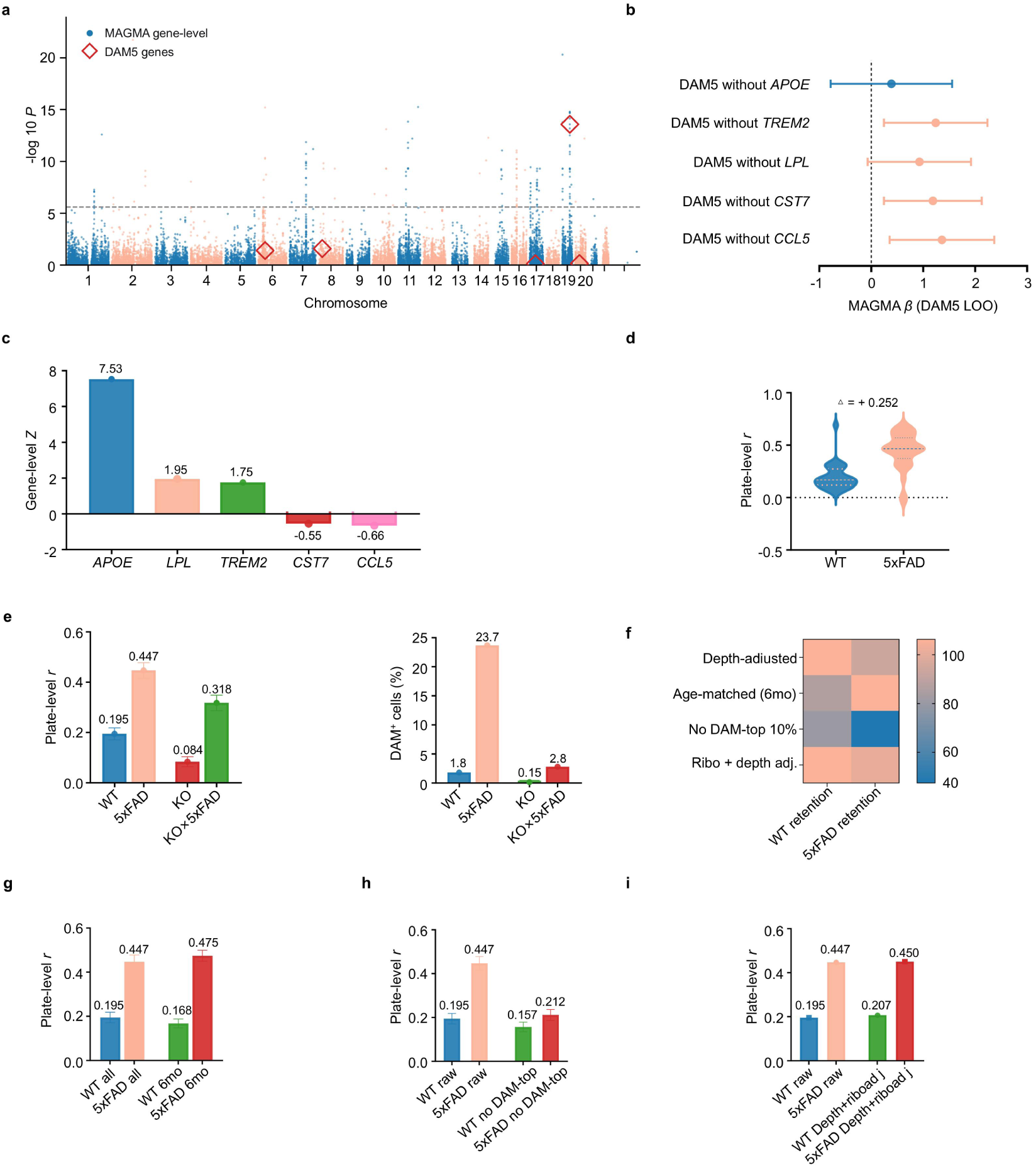
MAGMA genetic risk enrichment and coupling amplification in the 5xFAD model. a, Manhattan plot of MAGMA gene-level results (dots represent −log_10_*P*, red diamonds highlight DAM5 signature genes, and the horizontal dashed line indicates the genome-wide significance threshold). **b,** Forest plot of MAGMA *β* coefficients from LOO analyses of the DAM5 signature (error bars represent 95% CIs). **c,** Gene-level *Z*-scores for the five DAM5 signature genes. **d,** Violin plot of plate-level coupling *r* between C57BL/6 wild-type controls (WT, *n* = 28 plates) and 5xFAD mice (*n* = 31 plates). **e,** Plate-level coupling *r* (left) and DAM^+^ cell proportions (%; right) across the four experimental genotypes. f, Heatmap showing the percentage retention (%) of WT and 5xFAD plate-level *r* across four sensitivity specifications. **g–i,** Comparisons of plate-level *r* between all-age and 6-month-matched cohorts (g), raw data versus excluding the top 10% highest DAM-scoring cells (h), and raw data versus joint adjustment for sequencing depth and ribosomal gene fraction (i). Data in e, g–i are presented as mean ± s.e.m.

Next, we evaluated the impact of aggressive amyloidosis (5xFAD) and Trem2 genetic perturbation on coupling in dataset GSE98969, treating sequencing plates as independent statistical units (5xFAD: 31 plates, 3,968 cells; C57BL/6 wild-type [WT] controls: 28 plates, 4,869 cells; *Trem2*−KO×5xFAD : 6 plates; *Trem2*−KO 3 plates; Fig. 6d–i and Supplementary Table 2). In plaque-free WT control mice, all 28 sequencing plates exhibited positive coupling (plate-level mean *r* = 0.195 ± 0.024, one-sample *t*(27) = 7.04, *P* = 1.4 *×* 10^-7^), reaffirming that baseline coupling is an intrinsic property of homeostatic brain tissue. In 5xFAD mice, coupling was markedly amplified to *r* = 0.447 ± 0.031(30/31 plates positive; net increase over WT of Δ = +0.252, plate-level Mann–Whitney test *P* < 10^-4^, Cohen’s *d* = 1.68; Fig. 6d).

Plate-level coupling across the four genotypes followed a clear monotonic staircase:*Trem2*−KO (0.084) < WT (0.195) < *Trem2−*KO×5xFAD (0.318) < 5xFAD (0.447; Fig. 6e, left). Crucially,in *Trem2*−KO×5xFAD double-mutant mice—despite *Trem2* deficiency causing the mature DAM^+^ cell subpopulation to collapse from 23.7% in 5xFAD to 2.8% (approaching the 1.8% seen in WT; Fig. 6e, right)—lysosomal–DAM coupling (0.318 ± 0.031, 6/6 plates positive, *P* = 2.0 *×* 10^-4^) remained significantly elevated above WT controls, retaining roughly half (Δ = +0.123) of the disease-induced amplification. Consistently, excluding the top 10% highest DAM-scoring cells in 5xFAD, restricting analyses to age-matched 6-month-old cohorts, or jointly adjusting for sequencing depth and ribosomal gene fraction preserved significant inter-group amplification (Fig. 6f–i and Supplementary Table 2). This genetic dissociation demonstrates that disease-induced lysosomal–DAM coupling amplification comprises two layers: a basal cell-intrinsic amplification driven directly by lysosomal burden independent of *TREM2*-mediated subpopulation expansion, and a secondary amplification contributed by *TREM2*–*APOE*-mediated expansion of mature DAM.

### Homeostatic maintenance of coupling across human AD Braak stages and a cross-system state-transition synchrony timescale model

Finally, leveraging the finely staged SEA-AD cohort (84 donors, median age 90 years, 236,002 microglial nuclei; Supplementary Table 4), we examined how lysosomal–DAM coupling evolves across the decades-long trajectory of chronic human neurodegeneration. At the donor-mean level, the lysosomal program and DAM score exhibited a strong positive correlation (*r* = 0.554, 95% CI: 0.385–0.686, *P* = 4.7 *×* 10^-8^; depth-adjusted partial *r* = 0.544, retaining 98.3%; Fig. 7a, b). Within individual donors (within-donor *r*), coupling was stably maintained at a high level throughout pathological progression from Early (Braak 0–II, *n* = 6, mean *r* = 0.264) and Mid (Braak III–IV, *n* = 29, *r* = 0.272) to Late stages (Braak V–VI, *n* = 49, *r* = 0.309; reaching 0.337 in Braak VI), with one-sample *t*-tests yielding *P* < 10^-3^ across all stages and an overall median (0.302) exceeding those of the other human cohorts (Fig. 7c**–**e).

**Fig. 7.**
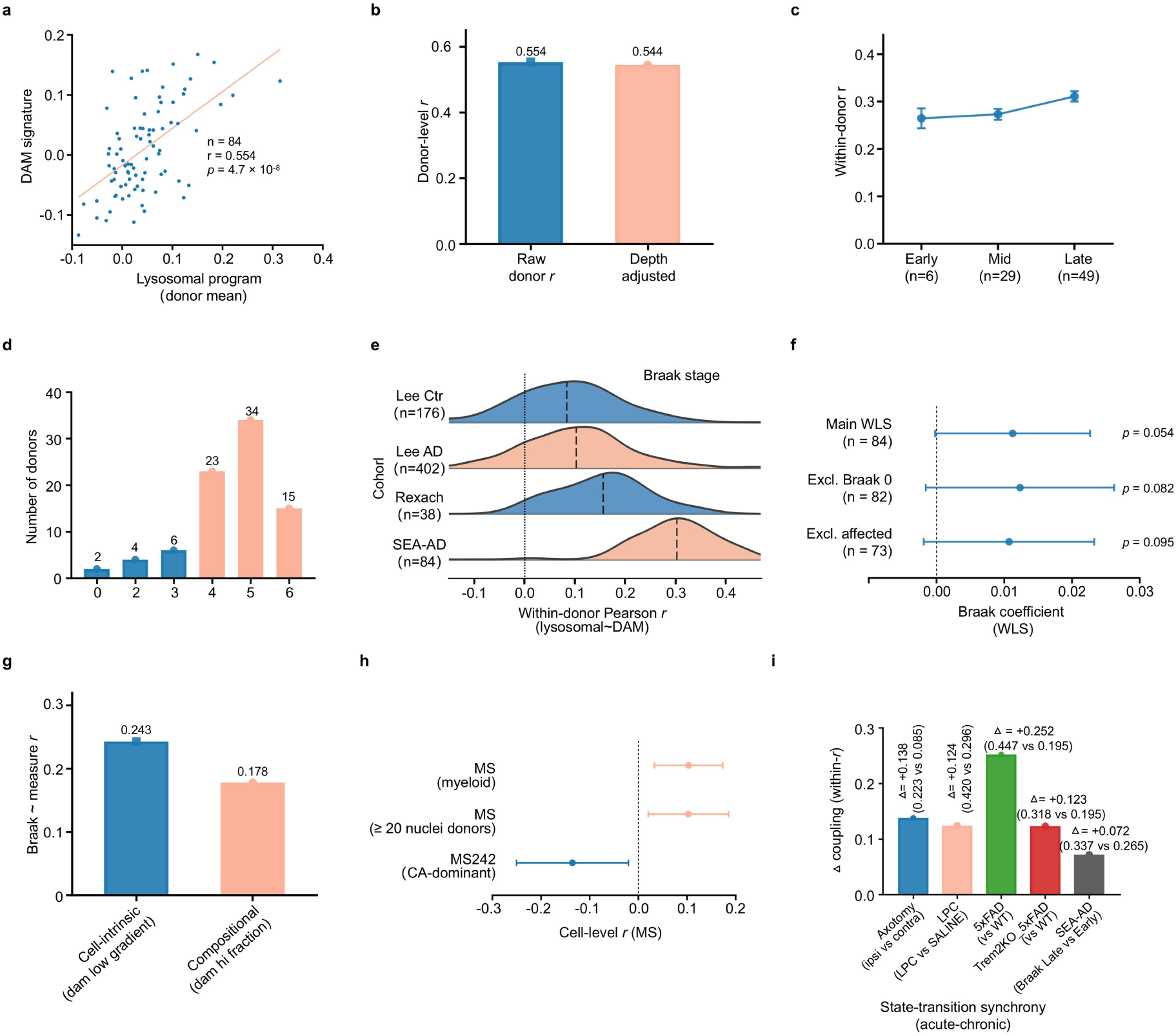
Lysosomal–DAM coupling and Braak stage dynamics in the SEA-AD cohort (84 independent donors, 236,002 microglial nuclei). **a**, Scatter plot of donor-mean lysosomal program scores versus DAM signature scores across all 84 donors. **b**, Comparison of raw versus sequencing depth-adjusted donor-level Pearson *r*. **c**, Within-donor correlation coefficients (within-donor *r*, mean ± s.e.m.) across coarse Braak stage bins: Early (stages 0–II, *n* = 6), Mid (stages III–IV, *n* = 29), and Late (stages V–VI, *n* = 49). **d**, Distribution of donor counts across individual Braak stages in the SEA-AD cohort (stage 0: *n* = 2; II: *n* = 4; III: *n* = 6; IV: *n* = 23; V: *n* = 34; VI: *n* = 15). **e**, Kernel density distributions of within-donor Pearson *r* across cohorts and diagnostic groups: Lee controls (*n* = 176), Lee AD (*n* = 402), Rexach (*n* = 38), and SEA-AD (*n* = 84); vertical dashed lines indicate group medians. **f**, Forest plot of WLS Braak regression coefficients (b) across the full cohort (*n* = 84), excluding Braak 0 donors (*n* = 82), and excluding severely affsected donors (*n* = 73; error bars represent 95% CIs). **g**, Comparison of Braak stage correlations for the cell-intrinsic component (residual gradient after excluding the top 10% DAM-high nuclei) versus the compositional component (fraction of DAM-high nuclei). **h**, Cell-level coupling sensitivity analysis in human multiple sclerosis (MS) overall myeloid cells, donors with ≥ 20 nuclei, and a chronic active (CA) lesion-dominant donor (MS242). **i**, Cross-system comparison of coupling magnitude shifts (Δ within-r) spanning acute murine insults and chronic human disease (see Table 1 for details).

**Table 1.** Summary of lysosomal–DAM coupling change magnitudes (Δ) across experimental models and human disease cohorts. Note: Δ is defined as the difference in coupling between the disease or injury state and the corresponding control state within the same dataset (difference in Fisher *z*-pooled *r*, or difference in mean plate-level or donor-level *r*). The 95% confidence intervals (CIs) for Δ and their calculation specifications are detailed in Supplementary Table 16.

| System (statistical unit) | Control state (*r*) | Disease or injury state (*r*) | Δ | 95% CI of Δ |
|---|---|---|---|---|
| Facial nerve axotomy (GSE90975, cell level) | Contralateral 0.085 | Ipsilateral 0.223 | +0.138 | −0.01 to +0.28 |
| LPC demyelination (GSE121654, cell level) | Saline 0.296 | LPC 0.420 | +0.124 | +0.06 to +0.19 |
| 5xFAD (GSE98969, plate level) | C57BL/6 0.195 | 5xFAD 0.447 | +0.252 | +0.17 to +0.34 |
| *Trem2*-KO ×<br>5xFAD (plate level) | C57BL/6 0.195 | *Trem2*-KO × 5xFAD 0.318 | +0.123 | +0.04 to +0.21 |
| SEA-AD (donor level) | Braak Early 0.264 | Braak Late 0.337 | +0.072 | Spans zero |

However, formal testing for stage-dependent amplification revealed that incremental coupling enhancement across chronic human AD progression is minimal (Fig. 7f). Weighted least-squares (WLS) regression adjusting for age, sex, and nuclei count showed only a marginal trend (*b* = 0.0112, *P* = 0.054) that attenuated further upon excluding either the 2 Braak 0 donors (*P* = 0.082) or the 11 severely affected donors (*P* = 0.095), and pairwise comparisons across coarse stage bins were non-significant after Bonferroni correction (Fig. 7f and Supplementary Table 4c). Compositional decomposition demonstrated that the stage-associated rise in DAM scoring was driven primarily by continuous cell-intrinsic upregulation after excluding the top 10% DAM-high nuclei (*r* = 0.243, *P* = 0.026) rather than proportional expansion of the DAM-high subpopulation (*r* = 0.178, *P* = 0.105; Fig. 7g). In an exploratory extension to human multiple sclerosis (MS) white matter demyelinating lesions (*n* = 5 donors), overall myeloid cells and the majority of individual donors similarly maintained positive coupling, with only a single donor dominated by chronic active lesions (MS242) exhibiting negative decoupling (Fig. 7h and Supplementary Fig. 16).

Integrating the coupling change magnitudes (Δ)across all five experimental and clinical systems revealed a striking scaling principle across pathological timescales (Fig. 1d, Fig. 7i, Table 1, and Supplementary Table 20): coupling amplification was largest in genetically driven aggressive murine amyloidosis (5xFAD Δ = +0.252), intermediate in acute synchronized murine neural injury and demyelination (facial nerve axotomy Δ = +0.138; LPC demyelination Δ = +0.124; *Trem2*−KO×5xFAD Δ = +0.123),and smallest in decades-long sporadic human AD (SEA-AD Late versus Early Braak stages Δ = +0.072), roughly one-third the magnitude of 5xFAD). This cross-scale contrast points toward a “state-transition synchrony” model: acute toxin lesions or aggressive multi-transgene overexpression trigger a temporally compressed, highly synchronized phagocytic wave that recruits a large fraction of microglia simultaneously into lysosomal stress and DAM conversion; conversely, sporadic human AD unfolds asynchronously over decades, such that a single post-mortem cross-sectional snapshot captures microglia dispersed across desynchronized phases of engulfment and transcriptional adaptation, manifesting as robust cell-intrinsic baseline maintenance rather than an explosive jump in population slope.

## Discussion

Microglial reprogramming toward the DAM state is a shared core response to CNS injury and neurodegeneration, yet the upstream intracellular transduction logic linking extracellular pathological burden to nuclear transcriptional remodeling has long lacked quantitative in vivo definition [1–4]. By integrating cross-species, cross-disease single-cell and single-nucleus transcriptomic datasets under rigorous independent statistical unit correction, our study establishes an evolutionarily conserved quantitative coupling between the MiT/TFE-regulated CLEAR lysosomal stress program and DAM reprogramming. Crucially, this coupling is not a binary switch turned on exclusively after injury; rather, it operates as an intrinsic basal homeostatic tone across uninjured, plaque-free brain tissue that undergoes context-dependent amplification during tissue damage and proteinopathy clearance.

The central conceptual advance of our work is the timescale model in which lysosomal–DAM coupling amplification scales with “state-transition synchrony” (Fig. 1d and Fig. 7i). Why do murine 5xFAD amyloidosis and acute demyelinating injury exhibit robust coupling amplification (Δ = +0.124–0.252), whereas aged human late-stage AD brain tissue (SEA-AD)—Bearing a massive cumulative pathological burden—displays only a modest stage-associated increase (Δ = +0.072)? From the perspective of clinical neuropathological kinetics, focal toxin injections or aggressive multi-transgene overexpression in 5xFAD mice generate a temporally compressed, highly synchronized wave of myelin debris or A*β* engulfment over days to months, forcing a large fraction of microglia to simultaneously experience lysosomal membrane tension surges and transcriptional reprogramming. Conversely, sporadic human AD unfolds over two to three decades, with neuronal degeneration and plaque deposition progressing asynchronously and focally across the parenchyma. In a single post-mortem cross-sectional snapshot, microglia are sampled across desynchronized phases of engulfment, lysosomal degradation, and transcriptional adaptation; consequently, their coupling signal manifests as high within-donor baseline maintenance and gradual cell-intrinsic upregulation rather than an explosive jump in population slope. While differences across species, sequencing modalities (single-cell versus single-nucleus RNA-seq), and gene-set mapping may also influence absolute effect sizes, state-transition synchrony provides a unifying mechanistic hypothesis for this cross-system gradient.

Dissecting the genetic and molecular architecture of this axis resolves the apparent conceptual tension between the conserved upstream role of the lysosomal program and the downstream localization of common AD genetic risk. MAGMA gene-set analysis demonstrated that common sporadic AD risk variants show no enrichment in the upstream lysosomal program (LYS10; *β* = 0.106, *P* = 0.357) but enrich strongly in the downstream DAM signature—a signal anchored almost entirely in *APOE* (*Z* = 7.53) rather than distributed polygenically across the module (Fig. 6a–c). Far from weakening an upstream pacemaking model, this architecture is evolutionarily and pathophysiologically coherent: core lysosomal biogenesis and ion/tension-sensing machinery (LYS10) represent essential cellular housekeeping infrastructure required for basal microglial survival and routine tissue surveillance, buffering them against common high-frequency polymorphisms via purifying selection—a view mirrored by the fact that rare, high-penetrance mutations in lysosomal regulators (such as *GRN* and *NPC1*) cause severe early-onset neurodegenerative syndromes rather than common late-onset AD. By contrast, late-onset sporadic AD genetic risk—exemplified by *APOE*—acts at the downstream effector bottleneck, governing the efficiency with which microglia translate an intact upstream lysosomal stress signal into lipid clearance and sustained DAM amplification.

Building upon this genetic architecture and our knockout perturbation data, we deconstruct lysosomal–DAM coupling into a two-component model. The first component is an evolutionarily conserved, cell-intrinsic baseline coupling: present in healthy tissue and robustly retained after removing *Apoe*/*APOE* or excluding the terminal repair subpopulation C1 (Fig. 4b and Supplementary Table 11), it converts routine phagolysosomal flux into basal transcriptional tone. The second component is an injury- or disease-induced amplification module: genetically anchored downstream by *APOE*, it is further potentiated during heavy lipid and protein aggregate clearance by *TREM2*-mediated expansion of mature DAM subpopulations [19]. Crucially, however, even when the mature DAM^+^ subpopulation is reduced by nearly 90% in *Trem2*-KO×5xFAD mice, roughly half of the disease-induced coupling amplification is preserved, demonstrating that signal transduction from upstream lysosomal stress to the DAM transcriptional program operates largely independently of *TREM2*-mediated population expansion. Together, these two lines of evidence converge on a dual-layer architecture: an evolutionarily conserved, *APOE*-independent, and non-strictly *TREM2*-dependent baseline coupling axis, overlaid by an *APOE*-anchored, *TREM2*-potentiated disease amplification module.

Furthermore, the transcriptional divergence within the MiT/TFE family observed in our large-scale human cohorts (*MITF* and *TFE3* mRNAs closely tracking DAM, whereas *TFEB* mRNA does not) carries important methodological and biological implications. Because *TFEB* acts primarily via dephosphorylation-dependent nuclear translocation [6–8], relying solely on single-cell *TFEB* transcript abundance readily yields false-negative conclusions; by computing the shared downstream CLEAR target module (LYS10) alongside *MITF*/*TFE3* transcript levels, our approach bridges biochemical mechanisms with transcriptomic readouts. This framework forms a tight evidentiary triangle with recent functional genetic studies: whereas Balak et al. [10] and Tejwani et al. ^11^ demonstrated that primary lysosomal storage defects drive maladaptive pro-inflammatory DAM phenotypes via *MITF*/*TFE3*, and Kim et al.[12] showed that *Tfeb* overexpression ameliorates 5xFAD pathology, our findings reveal that physiological phagocytic load during acute neural injury engages the very same lysosomal–MiT/TFE–DAM axis to coordinate an *Apoe*^-^ and innate-immunity-enriched tissue repair program. Thus, whether activating this shared lysosomal sensing axis leads to adaptive tissue repair or chronic neurodegenerative inflammation is dictated by the duration and clearance tractability of the upstream trigger.

Several limitations define the boundaries of this study (systematically graded in Fig. 1e and Supplementary Table 21). First, our reanalysis is strictly observational and transcriptomic. Although detectable expression of mechanosensitive and calcium channels (*Tmem63a* and *Mcoln1*) and the earlier population-level peak of the lysosomal program (day 7) relative to the DAM peak (day 30) establish temporal precedence and tight quantitative coupling, they do not constitute definitive causal proof of unidirectional mechanical driving or live single-cell lineage progression. Specifically, static snRNA-seq/scRNA-seq snapshots capture steady-state and disease-adapted transcriptional equilibria—reflecting a homeostatic “pacemaking” responsiveness—rather than real-time biophysical membrane tension, transient calcium fluxes, or rapid post-translational nuclear shuttling. Consequently, transcriptomic covariation alone cannot fully exclude parallel co-activation by shared upstream microenvironmental cues. Establishing a closed causal loop will require future in vivo loss- and gain-of-function interventions—such as microglia-specific conditional knockout or pharmacological blockade of upstream mechanosensitive channels (*Tmem63a* and *Mcoln1*) or MiT/TFE factors (*Mitf*, *Tfe3*, and *Tfeb*) across acute neural injury (axotomy or LPC demyelination) and chronic amyloidosis models—paired with live-cell fluorescent tension biosensors to directly test whether disrupting lysosomal stress sensing blunts DAM reprogramming. Second, the number of independent animals in the murine cohorts (3–4 mice per group) and the scarcity of early asymptomatic donors in the SEA-AD cohort (only 6 donors in Braak 0–II) limit the statistical power of animal-level rank tests and early human stage slope analyses. Based on our framework, we propose three directly testable predictions for future investigation: (i) conditional ablation of *Tmem63a*/*Mcoln1* or *Mitf*/*Tfe3* in microglia will markedly blunt DAM reprogramming during the synchronized window of acute neural injury; (ii) precisely controlling the temporal synchrony of phagocytic substrate delivery in vitro will quantitatively modulate the lysosomal–DAM coupling slope; and (iii) in longitudinal human cerebrospinal fluid biomarker cohorts, lysosomal–DAM coupling activity will preferentially track dynamic markers of acute axonal injury and synaptic turnover rather than static cumulative amyloid burden.

## Conclusions

In conclusion, this study establishes lysosomal stress sensing and DAM reprogramming as an evolutionarily conserved, *APOE*-anchored state-transition module that scales with pathological timescales, providing a quantitative theoretical roadmap for precisely modulating CNS injury repair and neurodegenerative disease.

## Abbreviations

Aβ: Amyloid-β
AD: Alzheimer’s disease
ADNI: Alzheimer’s Disease Neuroimaging Initiative
APOE: Apolipoprotein E
CI: Confidence interval
CLEAR: Coordinated Lysosomal Expression and Regulation
CNS: Central nervous system
DAM: Disease-associated microglia
DAM4: 4-gene core DAM score (excluding Apoe)
DAM5: 5-gene core DAM score
ERCC: External RNA Controls Consortium
FACS: Fluorescence-activated cell sorting
GEO: Gene Expression Omnibus
GO: Gene Ontology
GWAS: Genome-wide association study
IQR: Interquartile range
KEGG: Kyoto Encyclopedia of Genes and Genomes
KO: Knockout
LOO: Leave-one-out
LPC: Lysophosphatidylcholine
LYS10: 10-gene lysosomal program score
MAGMA: Multi-marker Analysis of GenoMic Annotation
MiT/TFE: Microphthalmia-associated transcription factor/TFE family
MPS-IIIA: Mucopolysaccharidosis IIIA
MS: Multiple sclerosis
MSSM: Mount Sinai School of Medicine
OLS: Ordinary least squares
PCA: Principal component analysis
PSP: Progressive supranuclear palsy
RADC: Rush Alzheimer’s Disease Center
REPAIR12: 12-gene injury-repair signature
scRNA-seq: Single-cell RNA sequencing
SD: Standard deviation
SEM: Standard error of the mean
SEA-AD: Seattle Alzheimer’s Disease Brain Cell Atlas
TFEB: Transcription factor EB
TFE3: Transcription factor binding to IGHM enhancer 3
TREM2: Triggering receptor expressed on myeloid cells 2
UMAP: Uniform Manifold Approximation and Projection
UMI: Unique molecular identifier
WLS: Weighted least squares
WT: Wild-type

## Author contributions

Y.L.: Conceptualization, Methodology, Formal analysis, Investigation, Writing – original draft, Writing – review & editing, Visualization. H.Y.: Formal analysis. Y.Y.Z.: Formal analysis. L.G.: Data curation. J.W.: Conceptualization, Supervision, Writing – review & editing. L.C.: Conceptualization, Supervision, Writing – review & editing. Y.H.Z.: Conceptualization, Supervision, Project administration, Writing – review & editing. All authors read and approved the final manuscript.

## Data availability

The data supporting the findings of this study are available within the article (Figs. 1–7, Table 1) and its supplementary materials (Supplementary Figs. 1–16 and Supplementary Tables 1–21). All single-cell and single-nucleus datasets reanalyzed in this study were previously published and are publicly available: the murine facial nerve axotomy dataset (GSE90975) [13], the murine corpus callosum LPC demyelination dataset (GSE121654) [2], the murine chronic amyloidosis dataset including 5xFAD and *Trem2*-knockout genotypes (GSE98969) [1], and the human multiple sclerosis white-matter cohort (GSE118257) [17] were obtained from the Gene Expression Omnibus (GEO, https://www.ncbi.nlm.nih.gov/geo/) . The SEA-AD cohort [16] was downloaded from the SEA-AD study website via the AD Knowledge Portal (Synapse accession syn26223298; https://adknowledgeportal.synapse.org/) . Standardized count matrices for the Lee 2026 cohort[14] and the Rexach 2024 cohort [15] 0were obtained from CZ CELLxGENE Discover (https://cellxgene.cziscience.com/collections/c53573b2-eff4-4c5e-9ad0-b24d422dfd9b, respectively). Alzheimer’s disease GWAS summary statistics were obtained from Bellenguez et al. (2022) via the GWAS Catalog (accession GCST90027158; https://www.ebi.ac.uk/gwas/studies/GCST90027158) . Gene set collections (KEGG_2019_Mouse, GO_Biological_Process_2023) are available from Enrichr (https://maayanlab.cloud/Enrichr/). The 1000 Genomes Project European phase 3 reference panel is available from the International Genome portal (https://www.internationalgenome.org/). Source data are provided with this paper.

## Code availability

All custom analysis scripts, configuration files, gene set definitions and intermediate results required to reproduce every figure and statistical test in this study have been publicly archived on Zenodo (Zenodo ((https://doi.org/10.5281/zenodo.10.5281/zenodo.23259475).

