## Supplementary Materials for "Evolutionarily conserved coupling of lysosomal response sensing and disease-associated microglial reprogramming scales with pathological timescales"

**Supplementary Methods | Mathematical Derivations, Statistical Formulas, and Implementation Benchmarks**

1. Ordinary least-squares (OLS) versus Liang–Zeger cluster-robust variance estimation in GSE90975

In the discovery cohort GSE90975 ($N$ = 1,437 cells across $G$ = 14 batch-by-timepoint-by-laterality strata), the single-cell linear interaction model is specified as:

$$y_i = \beta_0 + \beta_1 L_i + \beta_2 S_i + \beta_3 (L_i \times S_i) + \gamma_1 I(Day_i = 7) + \gamma_2 I(Day_i = 30) + \varepsilon_i \qquad (1)$$

with $K$ = 6 parameters. Naive OLS and cluster-robust OLS share the identical design matrix X and residual vector $\hat{\boldsymbol{\varepsilon}}$, yielding the identical point estimate $\hat{\beta}_3$=+0.1766. Under naive OLS (assuming independent and identically distributed homoscedastic errors), the variance–covariance matrix is:

$$\widehat{Var}_{OLS}(\hat{\boldsymbol{\beta}}) = \hat{\sigma}^2 (\mathbf{X}^\top \mathbf{X})^{-1}, \quad \hat{\sigma}^2 = \frac{1}{N-K} \sum_{i=1}^{N} \hat{\varepsilon}_i^2 \qquad (2)$$

yielding $SE_{OLS}(\hat{\beta}_3)$ = 0.0542, Wald statistic $W$=0.1766/0.0542=3.26, and two-sided $P$ = 0.00116.Under the Liang–Zeger sandwich estimator clustered by $g$=1,…,$G$, the cluster-robust variance–covariance matrix is:

$$\widehat{Var}_{CRSE}(\hat{\beta}) = c\,(\mathrm{X}^\top \mathrm{X})^{-1} \left( \sum_{g=1}^{G} \mathrm{X}_g^\top \ \hat{\varepsilon}_g \hat{\varepsilon}_g^\top \ \mathrm{X}_g \right) (\mathrm{X}^\top \mathrm{X})^{-1} \qquad (3)$$

where the finite-sample correction factor $c$ is given by:

$$c = \frac{G}{G-1} \cdot \frac{N-1}{N-K} = \frac{14}{13} \times \frac{1{,}436}{1{,}431} \approx 1.0807, \quad \sqrt{c} \approx 1.0396 \qquad (4)$$

This yields $SE_{CRSE}(\hat{\beta}_3)$=0.0590, Wald statistic $W$=0.1766/0.0590=2.99, two-sided asymptotic normal $P$=2(1−Φ(|$W$|))=0.00278, and 95% CI \$=0.1766±1.96×0.0590=[0.0609,0.2922]. The 8.9% inflation in standard error (0.0590/0.0542=1.089) decomposes into ~ 4.0% from the finite-sample multiplier $\sqrt{c}$\$*and*\$~ 4.9% from within-stratum cellular correlation.

2. Within-unit Pearson correlation, Fisher $z$-pooling, Cohen's $d$, and E-value transformations

Within each stratum, plate, or donor $k$ of sample size $n_k$, the single-cell Pearson correlation between lysosomal ($x$) and DAM ($y$ scores is:

$$r_k = \frac{\sum_{i=1}^{n_k}(x_i - \bar{x})(y_i - \bar{y})}{\sqrt{\sum_{i=1}^{n_k}(x_i - \bar{x})^2}\sqrt{\sum_{i=1}^{n_k}(y_i - \bar{y})^2}} \quad (5)$$

Each correlation $r_k$ is mapped to Fisher's $z$-scale via:

$$z_k = artanh(r_k) = \frac{1}{2}\ln\left(\frac{1+r_k}{1-r_k}\right) \quad (6)$$

and pooled across $K$ independent units using inverse-variance weights $w_k = n_k - 3$ (with $w_k = n_k$ and $w_k = 1$ evaluated in sensitivity analyses; Supplementary Fig. 6):

$$\bar{z} = \frac{\sum_{k=1}^{K} w_k z_k}{\sum_{k=1}^{K} w_k}, \quad SE(\bar{z}) = \frac{1}{\sqrt{\sum_{k=1}^{K}(n_k - 3)}} \quad (7)$$

The pooled correlation and its 95% confidence interval are obtained via the inverse hyperbolic tangent:

$$\bar{r} = \tanh(\bar{z}) = \frac{e^{2\bar{z}} - 1}{e^{2\bar{z}} + 1}, \quad CI_{95\%}(\bar{r}) = \tanh(\bar{z} \pm 1.96\, SE(\bar{z})) \quad (8)$$

For cross-dataset repair signature projections (GSE121654), standardized cell-level effect sizes between LPC ($n_1$) and saline ($n_2$) groups are computed as Cohen's $d$:

$$d = \frac{\bar{X}_{LPC} - \bar{X}_{Saline}}{s_{pooled}}, \quad s_{pooled} = \sqrt{\frac{(n_1 - 1)s_1^2 + (n_2 - 1)s_2^2}{n_1 + n_2 - 2}} \quad (9)$$

For E-value sensitivity analyses of unmeasured confounding (Supplementary Table 7b), continuous correlation coefficients $r$ are first converted to standardized mean differences $d = 2r/\sqrt{1 - r^2}$ (Chinn, 2000), transformed to approximate risk ratios via $RR \approx exp(0.91 \times d)$, and mapped to the E-value scale (VanderWeele & Ding, 2017):

$$E - value = RR + \sqrt{RR(RR - 1)} \quad (10)$$

3. Memory-efficient chunked score_genes benchmark in SEA-AD and UMAP spatial autocorrelation

In the SEA-AD cohort (236,002 nuclei × 36,601 genes), to bypass memory bottlenecks without altering scoring mathematics, we implemented a chunked sparse CSR matrix scanner replicating Scanpy's score_genes (n_bins = 25, ctrl_size = 50, random_state = 42). Across a 3-donor validation subset (H21.33.023, H21.33.043, H21.33.044; 5,675 nuclei), single-cell scores from the chunked implementation correlated with standard in-memory Scanpy score_genes at $r = 0.984$ (lysosomal program) and $r = 0.990$ (DAM signature; Supplementary Table 20c). Conversely, an exploratory gene-wise $z$-score mean implementation compressed within-donor correlations by 0.069–0.139 across the validation donors (within-r shifting from 0.358 to 0.231, 0.188 to 0.129, and 0.439 to 0.324; Supplementary Table 20b, c) and was therefore discarded from primary reporting.

To test for local spatial autocorrelation of module scores on UMAP manifolds in the Lee 2026 cohort, we constructed $k$-nearest-neighbor graphs ($k \in \{6,12,24\}$) on UMAP coordinates and computed Moran's $I$. Unadjusted Moran's $I$ ranged from 0.03 to 0.05 and decreased to ≈0 after residualizing scores on donor identity and sequencing depth across all values of $k$, confirming that visual clustering on UMAP embeddings reflects inter-donor baseline differences rather than intrinsic manifold autocorrelation.

### Results

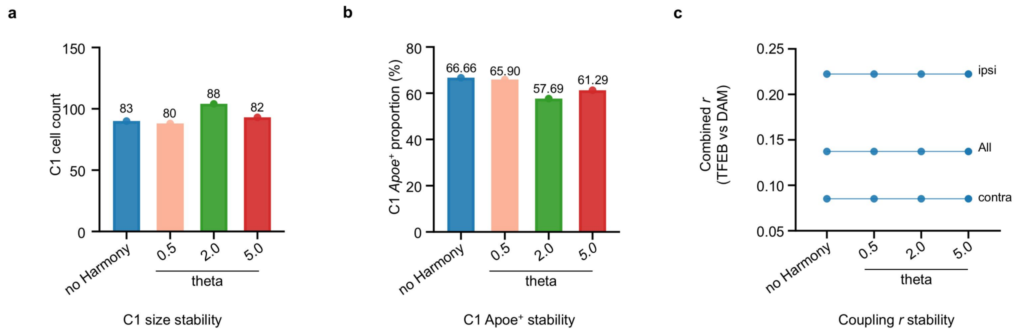


**Supplementary Fig. S1 | Robustness of the C1 subpopulation and coupling estimates across Harmony correction-strength parameter scans. a**, Number of C1 cells under each setting (no correction, and theta = 0.5, 2, and 5). **b**, Proportion of *Apoe*$^{+}$cells within C1 under each setting. **c,** Pooled coupling $r$ (contralateral, overall, and ipsilateral) under each setting.

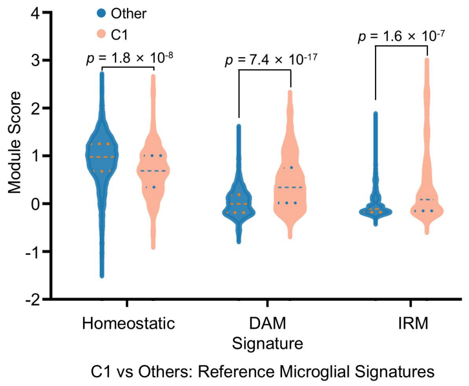


**Supplementary Fig. S2 | Annotation validation of the C1 subpopulation against reference microglial state signatures.** Module scores of three published reference signatures (homeostatic, DAM, and IRM) were computed for C1 ($n$ = 104) and the remaining clusters ($n$ = 1,333) and shown as box plots.

**a**

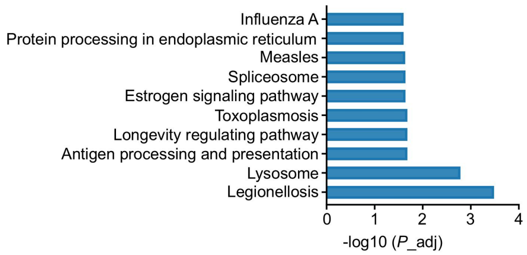


**b**

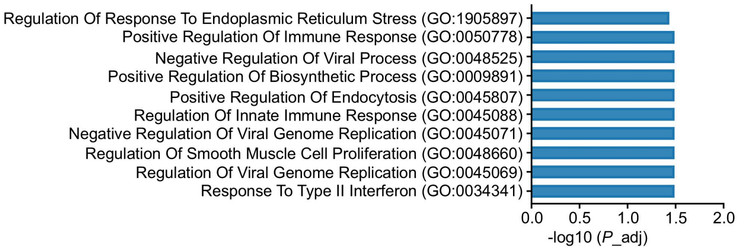


**c**

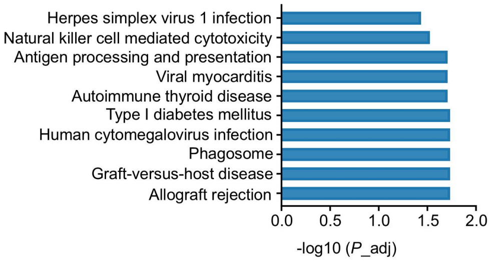


**Supplementary Fig. 3 | KEGG and GO biological process enrichment analyses of the stress-resolving signature. a**, Bar plot of KEGG pathway enrichment for the stress-resolving signature based on a local frozen snapshot. **b**, Bar plot of GO biological process enrichment for the stress-resolving signature. **c**, Bar plot of KEGG pathway enrichment for the repair signature after removing MHC-I genes (*H2-K1* and *H2-D1*).

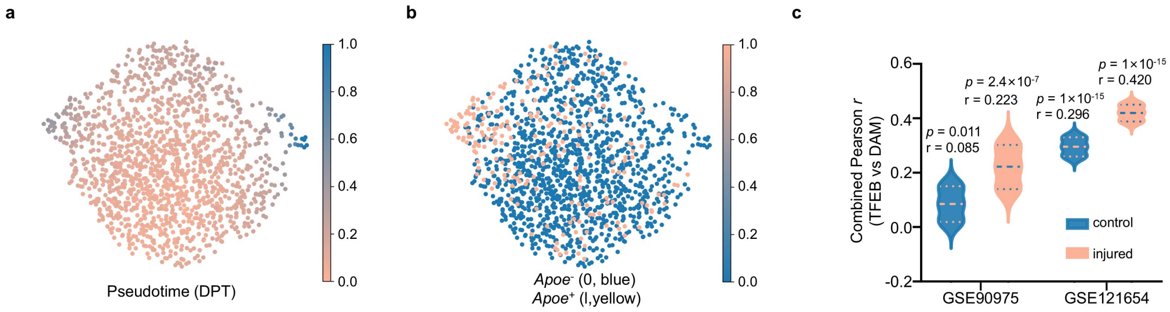


**Supplementary Fig. S4 | Transcriptional manifold and *Apoe*$^+$ cell identity in GSE90975. a,** Diffusion pseudotime (DPT) projected onto UMAP. **b,** UMAP distribution of *Apoe*$^-$ (blue) and *Apoe*$^+$ (yellow) cells. **c,** Pooled coupling *r* under control versus injury conditions across the two datasets.

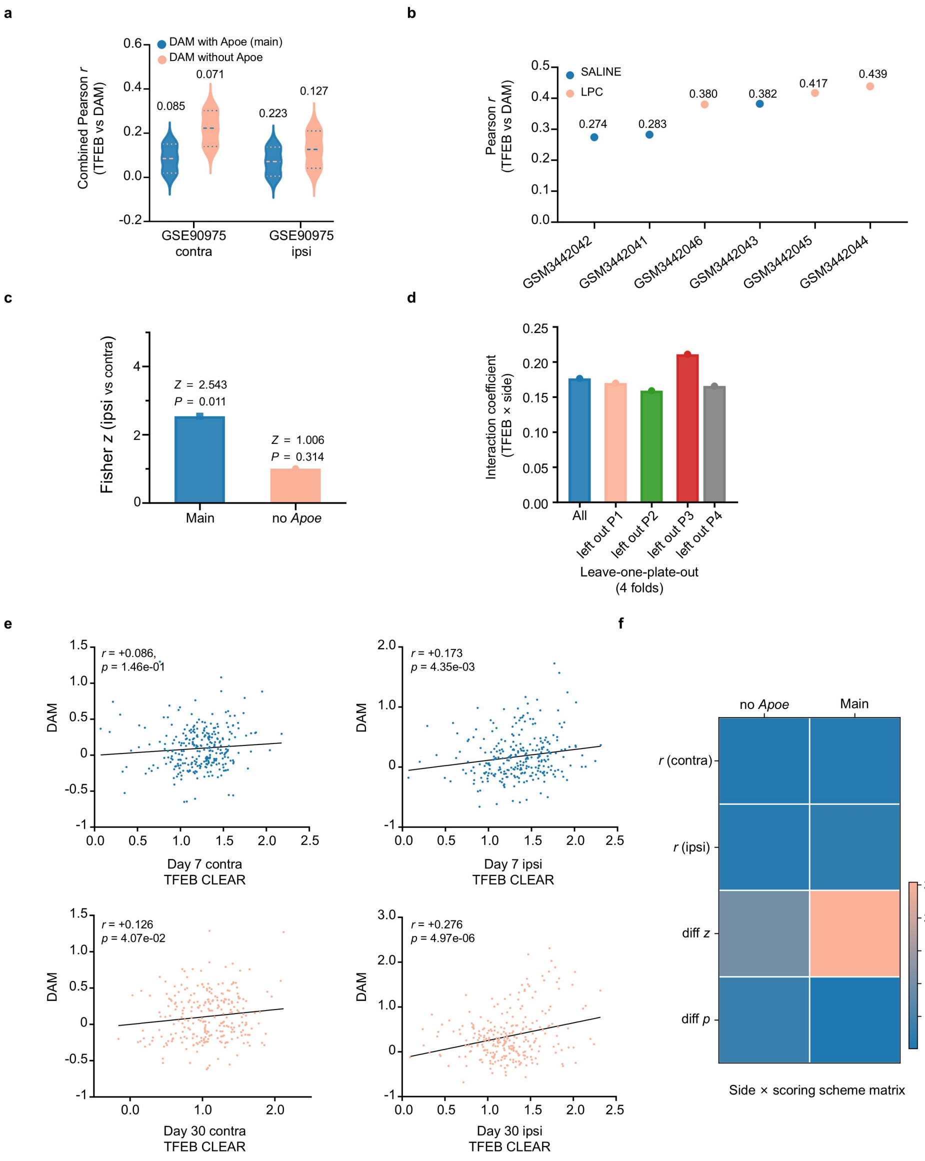


**Supplementary Fig. 5| Sensitivity analysis of *Apoe* removal, leave-one-plate-out cross-validation, and timepoint-stratified coupling scatter plots in murine injury models. a**, Within-group correlation distributions and pooled Pearson $r$ values (numbers at top) on the contralateral (contra) and ipsilateral (ipsi) sides of GSE90975 under the full DAM signature (blue) and the *Apoe*-removed DAM signature (orange). **b**, Dot plot of the six independent animals in GSE121654 (saline control in blue, LPC in orange) ordered by increasing within-animal single-cell Pearson $r$. **c**, Fisher $z$-statistics and $P$ values comparing ipsilateral versus contralateral coupling strength under the full (Main) and *Apoe*-removed (noApoe) signatures in GSE90975. **d**, Lysosomal program-by-laterality interaction coefficients in the full GSE90975 dataset (All) and across four-fold leave-one-plate-out cross-validation (sequentially leaving out P1–P4). **e**, Single-cell scatter plots and linear fits of lysosomal program (TFEB CLEAR) scores versus DAM scores on the contralateral (blue) and ipsilateral (orange) sides at days 7 and 30 in GSE90975. **f**, Heatmap matrix of correlation coefficients and between-side difference test statistics across laterality-by-scoring schemes (Main versus noApoe) in GSE90975.

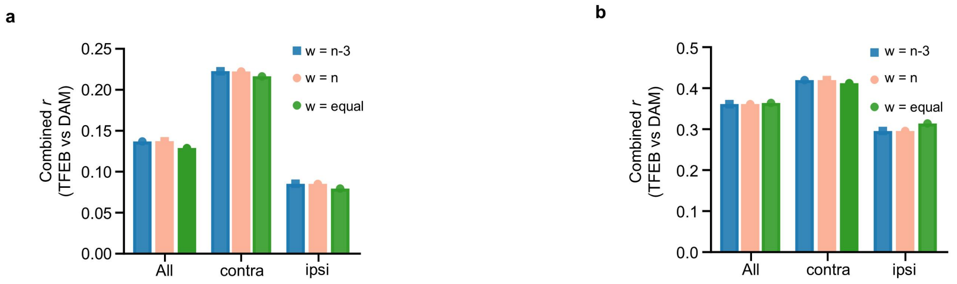


**Supplementary Fig. S6 Sensitivity of Fisher z pooling to weighting schemes.** A, Pooled *r* (overall, contralateral, and ipsilateral) in GSE90975 under w = *n*−3, w = *n*, and equal weighting. B, The same comparison in GSE121654.

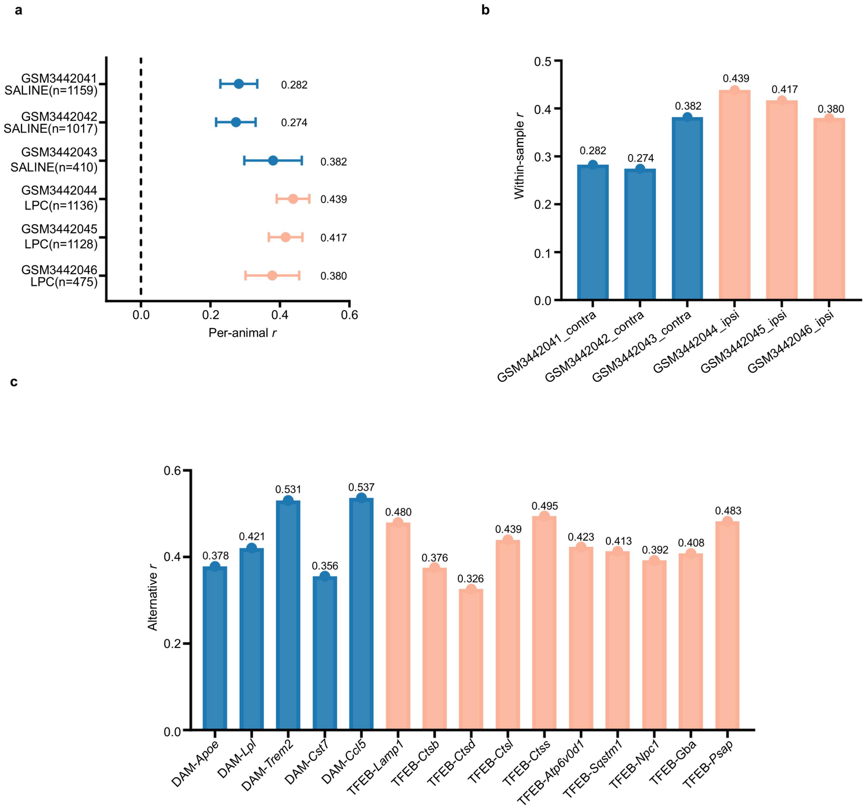


**Supplementary Fig. S7 | Per-animal and alternative-scoring sensitivity in GSE121654.** a, Per-animal *r* (95% CI) of the six animals. b, Within-sample-group r for each sample. c, Pooled r under alternative scoring and gene-subset specifications.

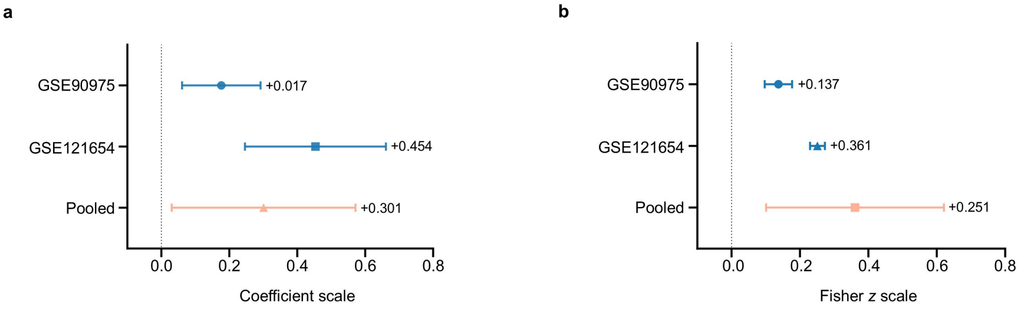


**Supplementary Fig. 8 | Random-effects meta-analysis of effect sizes across the two datasets.** a, Coefficient scale (GSE90975 +0.177, GSE121654 +0.454, pooled +0.301). b, Fisher *z* scale (GSE90975 +0.137, GSE121654 +0.361, pooled +0.251). Error bars denote 95% CI.

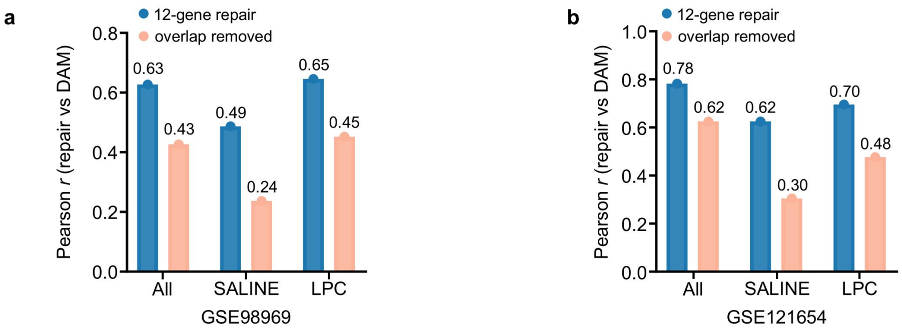


**Supplementary Fig. 9 | Shared-gene attenuation of repair-signature–DAM correlations at the cell level.**

a, b, Pearson correlation between the 12-gene repair signature and the DAM score before (blue) versus after (orange) removing the two shared genes (*Apoe*, *Ccl5*), computed at the cell level and stratified by condition.

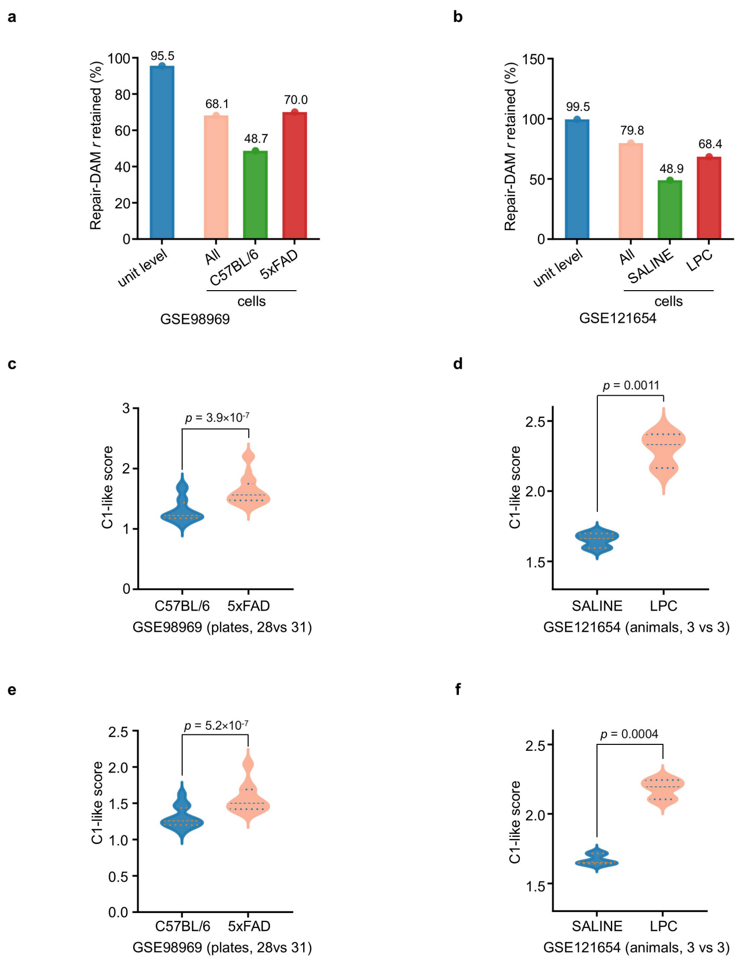


**Supplementary Fig. 10 | Unit-level projection of the C1-like repair state in external datasets and shared-gene attenuation of repair–DAM covariation. a, b**, C1 top-marker module scores (15 genes) at the inference unit. **c, d**, De-overlapped module (removing the DAM-core markers *Apoe* and *Cst7*). **e, f**, Retention rate of the repair-signature–DAM Pearson correlation after removing the two shared genes (*Apoe*, *Ccl5*; r_de-overlapped / *r*_full), comparing the inference-unit level with cell-level estimates stratified by condition.

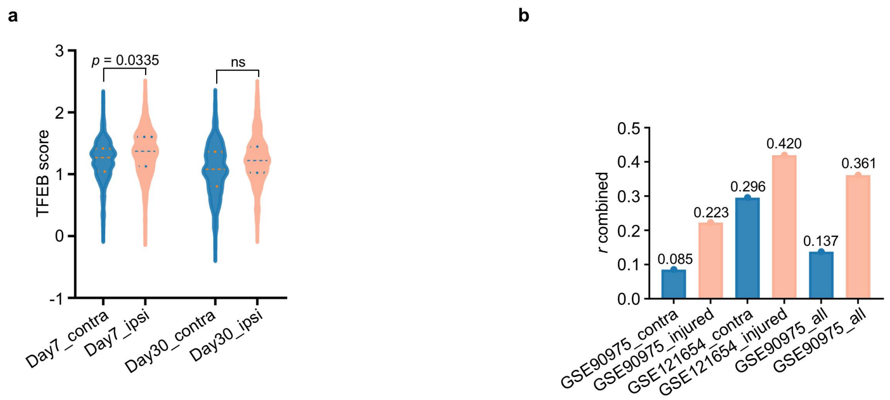


**Supplementary Fig. 11| Distribution of TFEB program scores. a,** Score distributions across timepoint-by-laterality groups. b, Pooled r across batch-by-condition combinations.

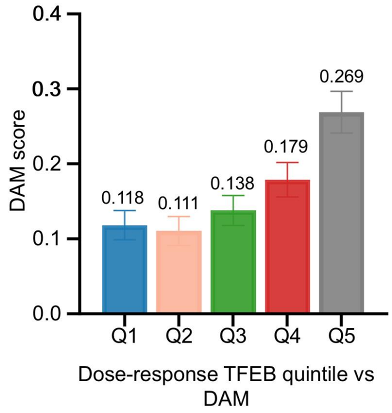


**Supplementary Fig. 12 | Quintiles of residualized TFEB scores versus DAM scores.** Mean DAM scores by Q1–Q5 of TFEB scores residualized on day × side (0.118, 0.111, 0.138, 0.179, and 0.269); error bars denote s.e.m.

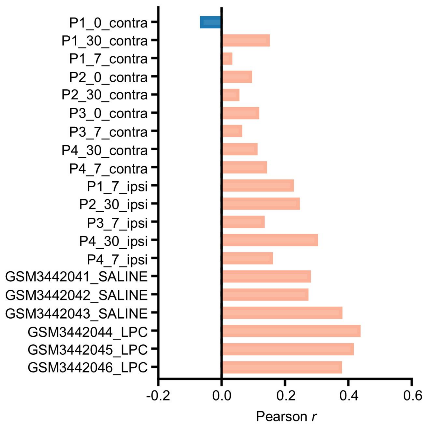


**Supplementary Fig. 13| Within-stratum correlation coefficients across all 20 batch-by-condition groups in the two murine injury datasets.** Horizontal bar plot showing within-group Pearson r values between the lysosomal program and the DAM signature across the 9 contralateral (contra) and 5 ipsilateral (ipsi) batch-by-timepoint strata in GSE90975 and the 6 individual animals (3 saline controls and 3 LPC-treated mice) in GSE121654.

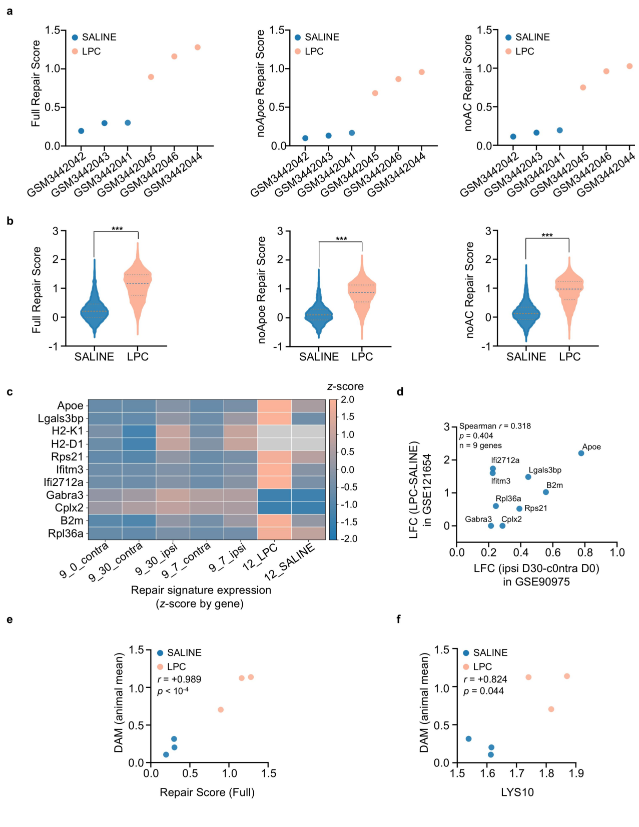


**Supplementary Fig. 14| Cross-model projection and validation of the 12-gene repair signature in the corpus callosum LPC demyelination dataset (GSE121654, *n* = 6 independent animals). a,** Animal-level mean repair scores across the three saline control (blue) and three LPC-treated (orange) mice under the full signature (Full), the Apoe-removed signature (noApoe), and the Apoe- and Ccl5-removed signature (noAC). **b,** Cell-level violin plots comparing repair scores between saline and LPC groups across the three signature versions $^{***}P < 0.001$. **c,** Heatmap of gene-wise z-score-standardized expression for repair signature genes across GSE90975 strata and GSE121654 LPC versus saline groups. **d,** Scatter plot of gene-level $\log_2$ fold changes (LFC) comparing day 30 ipsilateral versus day 0 contralateral microglia in GSE90975 against LPC versus saline microglia in GSE121654 across $n = 9$ mapped genes (Spearman $r = 0.318$, $P = 0.404$). **e,** Animal-level correlation between the full repair signature score and the mean DAM score across the six mice (Pearson $r = +0.989$, $P < 10^{-4}$). **f,** Animal-level correlation between the 10-gene lysosomal program score (LYS10) and the mean DAM score across the six mice.

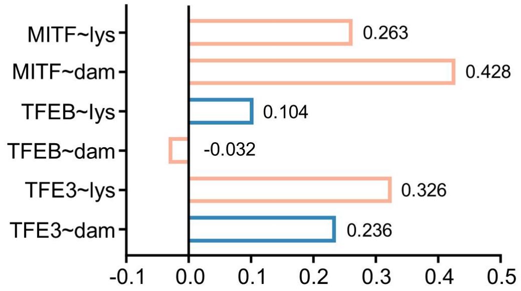


**Supplementary Fig. 15 | Raw correlations of MiT/TFE family transcripts in the Lee 2026 cohort.** Raw r (depth-unadjusted) of *MITF*, *TFEB* and *TFE3* transcripts against the lysosomal and DAM scores; depth-adjusted partial correlations are shown in Fig. 5c and Supplementary Table S13.

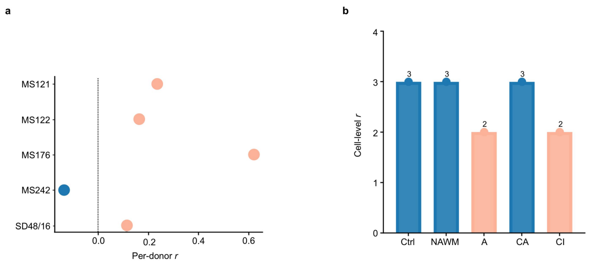


**Supplementary Fig. 16 | Donor-level and lesion-type views of coupling in the multiple sclerosis (MS) dataset.** a, Within-donor correlations for the five donors (MS121, MS122, MS176, MS242, and SD48/16). b, Cell-level coupling strength by lesion type (control white matter, normal-appearing white matter [NAWM], active, chronic active, and chronic inactive).

**Supplementary Table S1 | Differentially expressed genes in C1 versus the remaining clusters (top 15)**

| Group | Gene | $log_2FC$ | *P* | Adjusted *P* | Detection rate in C1 (%) | Detection rate in remaining clusters (%) |
|---|---|---|---|---|---|---|
| DAM core | *Apoe* | 4.36 | $3.80 \times 10^{-17}$ | $4.52 \times 10^{-14}$ | 57.7 | 11.7 |
| lysosomal | *Ctsl* | 1.78 | $3.15 \times 10^{-21}$ | $5.88 \times 10^{-18}$ | 91.3 | 67.5 |
| lysosomal | *Ctsb* | 1.40 | $8.67 \times 10^{-23}$ | $1.89 \times 10^{-19}$ | 97.1 | 85.4 |
| lysosomal | *Ctss* | 1.10 | $4.15 \times 10^{-29}$ | $1.81 \times 10^{-25}$ | 100 | 97 |
| lysosomal | *Psap* | 0.93 | $6.62 \times 10^{-10}$ | $1.44 \times 10^{-7}$ | 100 | 88.7 |
| lysosomal | *Lamp1* | 0.78 | $4.97 \times 10^{-5}$ | $4.43 \times 10^{-3}$ | 88.5 | 73.1 |
| lysosomal | *Ctsd* | 0.67 | $5.24 \times 10^{-13}$ | $2.54 \times 10^{-10}$ | 99 | 96.8 |
| MHC-I | *H2-D1* | 2.57 | $1.02 \times 10^{-35}$ | $1.34 \times 10^{-31}$ | 98.1 | 69.6 |
| MHC-I | *H2-K1* | 2.24 | $4.96 \times 10^{-23}$ | $1.30 \times 10^{-19}$ | 88.5 | 54.3 |
| MHC-I | *B2m* | 1.55 | $3.02 \times 10^{-34}$ | $1.98 \times 10^{-30}$ | 100 | 92.1 |
| MHC-I | *H2-T23* | 1.17 | $4.02 \times 10^{-6}$ | $4.42 \times 10^{-4}$ | 63.5 | 40.4 |
| other | *C4b* | 5.17 | $2.01 \times 10^{-7}$ | $2.86 \times 10^{-5}$ | 31.7 | 1.1 |
| other | *Ifitm3* | 4.24 | $1.71 \times 10^{-11}$ | $5.87 \times 10^{-9}$ | 43.3 | 4.4 |
| other | *Irf7* | 3.99 | $7.93 \times 10^{-7}$ | $1.01 \times 10^{-4}$ | 31.7 | 2.8 |
| other | *Fgl2* | 3.13 | $1.08 \times 10^{-6}$ | $1.35 \times 10^{-4}$ | 33.7 | 5.3 |

**Note:** Differential expression compared C1 (104 cells; Cluster 3) with the pooled remaining clusters (1,333 cells). *P* denotes two-sided unadjusted P-values; adjusted P denotes Benjamini–Hochberg-corrected values; detection rate denotes the percentage of cells with non-zero expression.

**Supplementary Table S2 | Pathway enrichment results for the two signatures**

| Signature | Database | Term | Adjusted *P* | Enriched genes | Interpretation |
|---|---|---|---|---|---|
| stress-resolving | KEGG | lysosomal | $1.6 \times 10^{-3}$ | CD68、GUSB、MAN2B1（3/11） | primary evidence |
| stress-resolving | KEGG | legionellosis | $3.3 \times 10^{-4}$ | ITGAM、HSPA1A、HSPA1B（3/11） | phagolysosome-related |
| stress-resolving | KEGG | antigen processing and presentation | $2.1 \times 10^{-2}$ | HSPA1A、HSPA1B（2/11） | supporting |
| stress-resolving | GO BP | regulation of TNF signaling | $2.9 \times 10^{-4}$ | GAS6、HSPA1A、HSPA1B（3/11） | supporting |
| stress-resolving | GO BP | regulation of dendritic cell chemotaxis | $2.9 \times 10^{-4}$ | GAS6、LGALS9（2/11） | supporting |
| stress- | GO BP | thermal | $2.9 \times 10^{-4}$ | HSPA1A、HSPA1B | hypothesis- |

| Signature | Database | Term | Adjusted *P* | Enriched genes | Interpretation |
|---|---|---|---|---|---|
| resolving | | adaptation | | （2/11） | generating |
| repair signature | GO BP | response to type II interferon | 3.2×10⁻² | IFITM3、CCL5（2/12） | primary evidence(non-MHC core) |
| repair signature | GO BP | negative regulation of viral genome replication | 3.2×10⁻² | IFITM3、CCL5（2/12） | interferon axis |
| repair signature | GO BP | regulation of innate immune response | 3.2×10⁻² | FCGR2B（1/12） | non-MHC core |

**Note:** Enrichment analyses used locally archived snapshots of KEGG 2019_Mouse and GO_Biological_Process_2023 (archived September 2026); terms were retained at Benjamini–Hochberg-adjusted $P < 0.05$. Owing to the small signature sizes, most terms are driven by 2–3 genes and should be interpreted as supporting annotations rather than unbiased pathway discoveries. KEGG enrichment of the repair signature was driven entirely by MHC-I genes; the five MHC-related terms share an identical gene set and adjusted P-value and are annotated once. Upon removing *H2-K1* and *H2-D1*, all significant repair-signature KEGG terms dropped to zero, whereas the three GO terms remained significant (adjusted $P = 3.2\times10^{-2}$), driven by *IFITM3*, *CCL5*, and *FCGR2B*.

**Supplementary Table S3 | Negative controls, conditional independence, and C1-exclusion analyses**

Panel a | Conditional independence (model: DAM ~ TFEB × is_C1 + side_ipsi + C(day); clustering $G = 14$)

| Item | Coefficient | 95%CI | *P* |
|---|---|---|---|
| TFEB main effect | +0.084 | +0.049 - +0.119 | $1.9\times10^{-6}$ |
| TFEB×C1 | +0.234 | −0.072 - +0.540 | 0.134 |

Panel b | C1-exclusion sensitivity

| Subset | n | Fisher *z r* | 95%CI | *P* | CRSEcoefficient | 95%CI | *P* |
|---|---|---|---|---|---|---|---|
| all cells | 1437 | +0.137 | +0.085 - +0.188 | $2.5\times10^{-7}$ | +0.177 | +0.061 - +0.292 | 0.0028 |
| C1 excluded | 1333 | +0.076 | +0.022 - +0.130 | 0.0061 | −0.012 | −0.106 - +0.083 | 0.810 |

Panel c | Negative-control outcomes

| Negative-control outcomes | Coefficient | 95%CI | *P* |
|---|---|---|---|
| ribosomal gene fraction | +0.560 | −0.053 - +1.173 | 0.074 |
| total UMI count | −302.46 | −630.92- +26.01 | 0.071 |
| cell-cycle score | +0.008 | −0.011 - +0.027 | 0.412 |

**Notes:** Panel b: Excluding C1 retained 55.5% of the overall correlation while fully abolishing the laterality interaction, paralleling the *Apoe*-removed specification (retained overall correlation, abolished laterality interaction) and indicating that C1 and *Apoe* jointly carry injury-induced amplification rather than baseline coupling. Panel c: The mitochondrial fraction was excluded as a negative control because of its direct mechanistic link to TFEB and mitophagy. 0/3 significant; no technical confounding signal detected.

**Supplementary Table S4 | Full robustness matrix of the two datasets**

| Analysis | GSE90975 coefficient | GSE90975 P | GSE121654 coefficient | GSE121654 P |
|---|---|---|---|---|
| cell-level OLS (reference) | 0.177 | 0.001 | 0.454 | $3.5\times10^{-17}$ |
| cell-level OLS, cluster-robust SE | 0.177 | 0.003 | 0.454 | 0.008[d] |
| mixed model, plate random intercept | 0.177 | 0.001 | not applicable[a] | not applicable |
| mixed model, animal random intercept | 0.177 | 0.001 | not applicable[a] | not applicable |
| mixed model, sample random intercept | not applicable | not applicable | 0.429 | $1.0\times10^{-16}$ |
| leave-one-plate-out (worst fold) | 0.159 | 0.012 | not applicable | not applicable |
| leave-one-sample-out (worst fold) | not applicable | not applicable | 0.325 | $4.1\times10^{-8}$ |
| pseudobulk sample-level regression † | −0.604 | 0.121 | 1.795 | 0.748 |
| rank-percentile scoring† | 0.059 | 0.151 | 0.188[b] | $6.2\times10^{-11}$ |
| Apoe-removed regression† | 0.029 | 0.577 | 0.387 | $1.0\times10^{-19}$ |
| random-effects meta-analysis | +0.301[b] | 0.029 | +0.251[b] | 0.032 |
| leave-one-animal-out, CRSE (worst fold) | not applicable | not applicable | +0.325 | 0.047，t(4) |
| exhaustive wild cluster bootstrap（$2^6$） | not applicable | not applicable | +0.454 | 0.094 |

**Note:** [a] Not performed in GSE121654. † Non-significant in the discovery cohort (1,437 cells), consistent with reduced statistical power; the corresponding tests are significant in the validation cohort. Pseudobulk sample-level regression probes between-sample effects rather than the cell-level coupling hypothesis and is included for methodological comparison; the two datasets are reported separately. Left column (0.301, 95% CI 0.022–0.456) and right column (0.251, 95% CI 0.031–0.572) are pooled Fisher z estimates; actual values are between-dataset pooled values, not single-dataset values. $P$ = 81%–90%; k = 2 is descriptive. Negative-control outcomes (ribosomal fraction, total UMI, cell-cycle score) were all non-significant ($P$ = 0.074, 0.071, 0.412); technical covariate adjustment (log UMI and log genes) retained 96% of the interaction (+0.169, $P$ = 0.004). $P$ values are two-sided and uncorrected. Pseudobulk regression (−0.604 and 1.795) reflects sample-level aggregation; after pooling, each batch–condition contributes only 1 observation (14 observations, $K$ = 6 in GSE90975), rendering the interaction estimate unstable with CIs crossing zero, a small-sample pooling artifact rather than directional counterevidence, disclosed for transparency. [d] t(5) conservative convention, clustering unit = animals ($G$ = 6); the coefficient is identical to cell-level OLS (+0.454), cluster-robust SE = 0.106, a 1.97-fold inflation over the naive SE; the animal-random-intercept mixed model gives +0.429, highly concordant with the CRSE.

**Supplementary Table S5 | Pooled coupling readouts via Fisher *z***

| No. | Analysis | | GSE90975 *r* | GSE90975 *P* | GSE121654 *r* | GSE121654 *P* |
|---|---|---|---|---|---|---|
| 1 | within-group correlation | control side | +0.085 | 0.011 | +0.296 | $<10^{-4}$[a] |
| 2 | | injury side | +0.223 | $2.4\times10^{-7}$ | +0.420 | $<10^{-4}$[a] |
| 3 | | overall | +0.137 | $2.5\times10^{-7}$ | +0.361 | $<10^{-4}$[a] |
| 4 | injury-versus-control side difference （Fisher *z*） | | not applicable | 0.011 | not applicable | $1.9\times10^{-7}$[b] |
| 4b | cell-level interaction term, cluster-robust SE | | not applicable | 0.003 | not applicable | $1.8\times10^{-5}$[b] |
| 5 | Apoe-removed, | control side | +0.071 | 0.034 | +0.244 | $<10^{-4}$[a] |
| 6 | | injury side | +0.127 | 0.004 | +0.337 | $<10^{-4}$[a] |
| 7 | | overall | +0.092 | $5.6\times10^{-4}$ | +0.293 | $<10^{-4}$[a] |
| 8 | | side difference （Fisher *z*） | not applicable | 0.314[c] | not applicable | $2.0\times10^{-4}$ |

**Note:** In GSE121654 the corresponding conditions are SALINE and LPC. Within-group Pearson correlations were computed per sample and pooled by cell-count-weighted Fisher z transformation. Between-group P-values were evaluated at the sample level by Fisher z tests, with independent biological samples as inferential units. *Apoe*-removed scores result from removing *Apoe* from the DAM signature. [a] Values reported as $<10^{-4}$ under double precision. [b] Row 4 cell-level side-difference tests were corroborated by animal-level cluster-robust interaction; the two are mutually consistent and interchangeable (Welch t (P = 0.081) and Mann–Whitney (P = 0.200) in the main text are equivalent tests; the two-sided P lower bound is 0.10 under 3-versus-3 design, reflecting limited power rather than imprecise testing). [c] The *Apoe*-removed increment in the discovery cohort was non-significant, consistent with the *Apoe*-removed regression result, attributable to limited cell numbers rather than effect absence. *Apoe*-removed sensitivity was cross-checked under three implementations: score_genes rs = 42 (0.244, 0.337, 0.293); score_genes rs = 0 (0.237, 0.338, 0.289); and deterministic z-mean implementation (0.254, 0.342, 0.300). Pooled r spans < 0.02 across implementations, with concordant direction and significance.

**Supplementary Table S6 | Meta-analysis and E-value summary**

Panel a | Random-effects meta-analysis

| Scale | Dataset | Estimate | 95%CI | *P* | Q （df=1） | P_Q | $\tau^2$ |
|---|---|---|---|---|---|---|---|
| coefficient scale | GSE90975 | +0.177 | +0.061 - +0.292 | 0.0028 | not applicable | not applicable | not applicable |
| coefficient scale | GSE121654 | +0.454 | +0.246 - +0.661 | 0.008[a] | not applicable | not applicable | not applicable |
| coefficient scale | pooled | +0.301 | +0.031 - +0.572 | 0.029 | 5.23 | 0.022 | 0.031 |
| Fisher z | GSE90975 | +0.137 | +0.085 - +0.188 | $2.5\times10^{-7}$ [c] | not applicable | not applicable | not applicable |
| Fisher z | GSE121654 | +0.361 | +0.260 - +0.450 | $<10^{-4}$ [c] | not applicable | not applicable | not applicable |
| Fisher z | pooled | +0.251 | +0.022 - +0.456 | 0.032 | 10.16 | 0.001 | 0.026 |

Notes (Panel a): [a] t(5) conservative convention; clustering unit = animals (G = 6). [b] Cell-level nominal P (pseudo-replication), reference only; for GSE121654 cluster-robust inference see the t(5) P = 0.008 note in the table. DerSimonian–Laird random-effects pooling; $k = 2$, an intrinsically low-information quantity. Fisher $z$, SE, $z$, Q, and $\tau^2$ are computed on the $z$ scale. The two studies' effects differ ~2.6-fold, compatible with the injury-model contrast. A sensitivity version with k extended to 4 (adding human donor-level estimates, donors as clustering units) exists; the main table retains k = 2 (mouse side only), with human results reported independently (Supplementary Tables S11, S12), to avoid mixing statistical-unit conventions.

Panel b | E-value analysis

| Dataset | *r* | ***RR*** **(approx)** | E-value (point) | E-value (CI end)[d] |
|---|---|---|---|---|
| GSE90975 (overall) | +0.137 | 1.29 | 1.89 | 1.61 |
| GSE90975(ipsilateral) | +0.222 | 1.51 | 2.40 | 1.91 |
| GSE121654 (overall) | +0.361 | 2.02 | 3.46 | 2.65 |
| GSE121654(LPC) | +0.420 | 2.32 | 4.07 | 3.74 |
| Lee 2026(donor level, AD) | +0.534 | 3.16 | 5.8 | 4.6 |
| Lee 2026(cell level) | +0.096 | 1.19 | 1.67 | not applicable |

**Notes (Panel b):** [d] Based on the CI end closest to the null. The entire table uses one conversion chain — for continuous exposure conventions: $r$ → standardized mean difference $d = 2r/\sqrt{(1-r^2)}$ → approximate $RR = \exp(0.91 \times d)$ → E-value $= RR + \sqrt{[RR(RR-1)]}$ (Chinn, 2000; VanderWeele & Ding, 2017). Examples: $r = 0.420$ → $d = 0.926$ → $RR = 2.32$ → E = 4.07; $r = 0.137$ → d = 0.277 → $RR = 1.29$ → E = 1.89; $r = 0.534$ → $d = 1.263$ → $RR = 3.16$ → E = 5.77 (reported as 5.8), with CI lower bound 0.460 → $d$ = 1.036 → $RR = 2.57$ → E = 4.57 (reported as 4.6); cell-level r = 0.096 → $d = 0.193$ → $RR = 1.19$ → E =

1.67. The E-value is the minimum confounding strength required to fully explain the observed association; mouse-side values (1.9–4.1) and the human value (5.8) should be read as quantitative references rather than precise thresholds.

**Supplementary Table S7| Decomposition, external gene sets, and technical covariates**

Panel a Association decomposition（*Apoe* as the carrier variable; cluster bootstrap with 1,000 resamples of the 14 clusters）

| Effect | Value | 95% CI | *P* |
|---|---|---|---|
| total association c | +0.107 | not applicable | $7.4\times10^{-7}$ |
| a-path (TFEB → *Apoe*) | +0.594 | not applicable | $7.9\times10^{-5}$ |
| b-path (*Apoe* → DAM4) | +0.044 | not applicable | $6.4\times10^{-7}$ |
| direct association c′ | +0.081 | not applicable | $6.6\times10^{-6}$ |
| indirect a×b | +0.026 | +0.005 - +0.044 | <0.001 |
| proportion via *Apoe* | 24.4% | not applicable | not applicable |

**Note:** a descriptive decomposition of association, not a causal mediation test; the term “mediation” is not used in the manuscript.

Panel b | Three-level correlation decomposition

| Level | r | *P* | Units | Description |
|---|---|---|---|---|
| pooled (all cells) | +0.171 | $6.5\times10^{-11}$ | 1437 | pseudo-replication, reference only[e] |
| within-group mean | +0.128 | not applicable | 14 | mean of the 14 within-group r values |
| between-group (14 means) | +0.450 | 0.107 | 14 | underpowered, descriptive only |

**Note:** [e]Pooled r = 0.171 is the single Pearson correlation across all cells; the main-text pooled r = +0.137 is the cell-count-weighted Fisher z pooling of 14 stratum correlations. The two are distinct statistics with no internal contradiction.

Panel c | External gene-set sensitivity

| Gene set | Genes used | Pooled r | 95%CI | *P* |
|---|---|---|---|---|
| GO_Lysosome | 23 | +0.085 | +0.033 - +0.137 | 0.0015 |
| Hammond_Repair | 13 | +0.253 | +0.203 - +0.301 | $<10^{-4}$ |
| lysosomal score (reference) | 10 | +0.137 | +0.085 - +0.188 | $<10^{-4}$ |
| KS2017_DAM [f] | 12 | +0.610 | +0.576 - +0.642 | $<10^{-4}$ |

Note: [f] Shares most component genes with the primary DAM signature (including *Apoe*、*Trem2*、*Lpl*、*Cst7*); the high correlation primarily reflects gene-set overlap and is not reported as independent coupling evidence.

Panel d | Technical covariate sensitivity

| Model | Coefficient | 95%CI | *P* | Retention |
|---|---|---|---|---|
| no covariates | +0.177 | +0.061 - +0.292 | 0.0028 | 100% |
| adjusted for log UMI | +0.169 | +0.055 - +0.283 | 0.0037 | 96% |
| adjusted for log genes | +0.169 | +0.056 - +0.283 | 0.0035 | 96% |
| all adjusted (incl. pct_mt) | +0.169 | +0.053 - +0.285 | 0.0044 | 96% |

**Note:** pct_mt may lie downstream of TFEB and mitophagy, so adjusting for it is a conservative potential over-adjustment; the 96% coefficient retention indicates the conclusion is insensitive to this choice.

**Supplementary Table S8 | Cross-dataset projection summary of the repair signature**

| Signature version | Signature genes | Detected genes[a] | Effective genes[b] | LPC mean | SALINE mean | Difference | Cohen *d*[c] | MW P[d] | Separation[e] |
|---|---|---|---|---|---|---|---|---|---|
| Full | 12 | 10 | 8 | +1.101 | +0.260 | +0.841 | 1.925 | 0.10 | 3/3 |
| noApoe | 11 | 9 | 7 | +0.828 | +0.135 | +0.693 | 1.846 | 0.10 | 3/3 |
| noAC | 10 | 8 | 6 | +0.902 | +0.158 | +0.743 | 1.861 | 0.10 | 3/3 |

Note: [a] Detected genes = signature genes present in the GSE121654 gene list (*H2-K1* and *H2-D1* absent). [b] Effective genes = non-zero expression in the pooled LPC/SALINE subset (*Gabra3* and *Cplx2* not expressed). [c] Cohen's *d* computed with the pooled within-group SD; providing a cell-count-independent effect-size metric, sample-level SDs are not meaningful and are not reported. [d] Sample-level Mann–Whitney U test; the two-sided mathematical lower bound of *P* is 0.10 for two fully separated groups of 3, to be read as a lower bound rather than non-significance. [e] Separation = number of LPC animals exceeding the maximum SALINE score; 3/3 = complete separation.

**Supplementary Table S9 | Variance decomposition of the DAM score (GSE90975, cell-level OLS**

| Model | $R^2$ | $\Delta R^2$ | Variance share |
|---|---|---|---|
| null model (intercept) | 0.000 | not applicable | not applicable |
| add Day | 0.017 | +0.017 | 1.73% |
| add Day and Side | 0.050 | +0.033 | 3.28% |
| add Day, Side, and TFEB | 0.074 | +0.024 | 2.37% |
| add Day, Side, TFEB, and TFEB × Side | 0.081 | +0.007 | 0.68% |

**Supplementary Table S10 | Within-group Pearson correlations across batch × timepoint × laterality strata in GSE90975**

| Stratum | Laterality | n cells | r (main score) | *P* | *r* (Apoe-removed) |
|---|---|---|---|---|---|
| P1_0_contra | contralateral | 82 | −0.068 | 0.546 | −0.029 |
| P1_30_contra | contralateral | 92 | 0.153 | 0.146 | 0.105 |
| P1_7_contra | contralateral | 96 | 0.034 | 0.742 | 0.109 |
| P2_0_contra | contralateral | 93 | 0.096 | 0.359 | 0.137 |
| P2_30_contra | contralateral | 79 | 0.057 | 0.619 | 0.072 |
| P3_0_contra | contralateral | 177 | 0.119 | 0.115 | 0.053 |
| P3_7_contra | contralateral | 94 | 0.065 | 0.536 | −0.107 |
| P4_30_contra | contralateral | 94 | 0.114 | 0.275 | 0.092 |
| P4_7_contra | contralateral | 94 | 0.144 | 0.166 | 0.211 |
| P1_7_ipsi | ipsilateral | 96 | 0.228 | 0.025 | 0.115 |
| P2_30_ipsi | ipsilateral | 173 | 0.247 | 0.001 | 0.146 |
| P3_7_ipsi | ipsilateral | 88 | 0.136 | 0.207 | 0.066 |
| P4_30_ipsi | ipsilateral | 93 | 0.304 | 0.003 | 0.184 |
| P4_7_ipsi | ipsilateral | 86 | 0.162 | 0.136 | 0.099 |

Note: 14 strata in total (batch × timepoint × laterality): 9 contralateral and 5 ipsilateral. The main score comprises *Apoe*, *Lpl*, *Trem2*, *Cst7* and *Ccl5*，the *Apoe*-removed version excludes *Apoe*. *P* denotes two-sided unadjusted values. Under the main score, 3 of 5 ipsilateral strata are significant; after removing *Apoe*, between-side groups remain significant, consistent with the side-difference analysis in Supplementary Table 5. Fisher z values can be derived from r via inverse transformation.

**Supplementary Table S11 | Mouse and human cell-level Fisher z-pooled correlation summary**

| Cohort | Group | Donors or animals | Cells or nuclei | *r* | 95%CI | *P* |
|---|---|---|---|---|---|---|
| Mouse GSE90975 | All | 10 | 1437 | +0.137 | +0.085 - +0.188 | $2.5\times10^{-7}$ |
| Mouse GSE121654 | All | 6 | 5325 | +0.361 | +0.260 - +0.450 | $<10^{-4}$ |
| mouse random-effects pooled | All | not applicable | not applicable | +0.251 | +0.022 - +0.456 | 0.032 |
| Lee 2026 | All | 578 | 100,379 | +0.096 | +0.090 - +0.102 | see main text |
| Lee 2026 | AD | 402 | not applicable | +0.098 | +0.091 - +0.106 | see main text |
| Lee 2026 | control | 176 | not applicable | +0.090 | +0.079 - | see main |

| Cohort | Group | Donors or animals | Cells or nuclei | *r* | 95%CI | *P* |
|---|---|---|---|---|---|---|
| | | | | | +0.102 | text |
| Rexach 2024 | All | 38 | 21,572 | +0.174 | +0.161 - +0.187 | see main text |
| Rexach 2024 | AD | 10 | not applicable | +0.203 | +0.181 - +0.225 | see main text |
| Rexach 2024 | control | 9 | not applicable | +0.099 | +0.066 - +0.131 | see main text |
| Rexach 2024 | PSP | 11 | not applicable | +0.150 | +0.124 - +0.176 | see main text |
| Rexach 2024 | Pick | 8 | not applicable | +0.209 | +0.183 - +0.235 | see main text |

Note: Human side: donor-stratified Fisher z pooling (w = $n$−3); mouse side follows the Supplementary Table 6 convention. Human cell-level 95% CIs are computed over total nuclei and are descriptive summaries under a pseudo-replication convention (precision overestimated), not for inference; human inference is anchored at the donor level (Supplementary Table 12). Per-donor correlation coefficients for Rexach are provided as an accompanying Supplementary Data table.

**Supplementary Table S12 | Human donor-level estimates and robustness matrix**

| Estimation | Cohort and group | Estimate | 95%CI |
|---|---|---|---|
| donor-mean correlation | Lee AD | +0.534 | +0.460 - +0.601 |
| donor-mean correlation | Lee control | +0.474 | +0.351 - +0.581 |
| donor-mean correlation | Lee all donors | +0.509 | +0.446 - +0.567 |
| donor-mean correlation | Lee MSSM subcohort | +0.527 | not applicable |
| donor-mean correlation | Lee RADC subcohort | +0.560 | not applicable |
| depth partial correlation | Lee | retained 101% | not applicable |
| APOE-removed, donor level | Lee AD | +0.267 | +0.174 - +0.356 |
| APOE-removed, cell level | Lee AD | +0.031 | not applicable |
| mixed-effects model (donor random intercept) | Rexach | +0.491（$P$=6.2×10$^{-12}$） | not applicable |
| depth + region partial correlation | Rexach | +0.532（107.5% retained） | +0.356 - +0.671 |
| donor × region unit level (descriptive only) | Rexach | +0.494 | not applicable |
| interaction lys × AD_flag | Lee（n=578） | +0.053（$P$=0.476） | not applicable |
| interaction lys × | Rexach（~19 donors | +0.432（$P$=0.008） | not applicable |

| Estimation | Cohort and group | Estimate | 95%CI |
|---|---|---|---|
| AD_flag | entering the model） | | |

Note: The Rexach donor × region units ($n$ = 82) are nested within 38 donors (pseudo-replication); unit-level correlations are descriptive only, and inference is anchored on the donor-random-intercept mixed model. The donor × region cross table is provided as an accompanying Supplementary Data table.

**Supplementary Table S13 | Donor-level partial correlations of MiT/TFE family transcripts with DAM and lysosomal scores (Lee 2026)**

| Transcript | Target score | Raw r | Depth-adjusted *r* | Detection-rate-adjusted *r* |
|---|---|---|---|---|
| MITF | DAM | N/A | +0.423（$P=1.7\times10^{-26}$） | +0.113 |
| TFE3 | DAM | N/A | +0.220（$P=9.6\times10^{-8}$） | +0.020 |
| TFEB | DAM | N/A | −0.114（$P=6.3\times10^{-3}$） | −0.129 |
| TFEB | lys | +0.104 | −0.163（$P=8.0\times10^{-5}$） | N/A |

Note: Partial correlations computed at the donor level; full values for the raw, depth-adjusted, depth + nGenes, and detection-rate-adjusted specifications are provided in the accompanying Supplementary Data. Detection rate is a partially endogenous covariate: if the association is real, DAM-enriched donors naturally show higher transcript detection rates, so the detection-rate-adjusted estimate should be interpreted as a conservative lower bound rather than a true effect.

**Supplementary Table S14 | MAGMA gene-set enrichment results and REPAIR12 mapping status**

**Panel a | MAGMA gene-set enrichment results (b37-aligned)**

| Gene set | NGENES | BETA | SE | *P* | Bonferroni |
|---|---|---|---|---|---|
| LYS10 | 10 | 0.106 | 0.290 | 0.357 | not significant |
| DAM5 | 5 | 1.947 | 0.453 | $8.7\times10^{-6}$ | significant |
| REPAIR8 | 8 | 0.439 | 0.300 | 0.071 | not significant |
| GO_LYSOSOME | 35 | 0.212 | 0.153 | 0.083 | not significant |
| GO_AUTOPHAGY | 98 | 0.030 | 0.093 | 0.372 | not significant |
| KEGG_AD (positive control) | 146 | 0.209 | 0.077 | 0.0032 | significant |

Note: Bonferroni threshold = 0.05/6 = 0.0083. Total genes analyzed = 18,349; mean sample size ≈ $4.7\times10^{5}$ (gene-level median 467,888; range 42,140–487,511). Gene sets were tested in a joint conditional model.

**Panel b | Leave-one-gene-out analysis of DAM5**

| Removed gene | Remaining genes | BETA | *P* |
|---|---|---|---|
| *APOE* | *TREM2*, *LPL*, *CST7*, *CCL5* | 0.389 | 0.257 |

| Removed gene | Remaining genes | BETA | *P* |
|---|---|---|---|
| *TREM2* | *APOE*, *LPL*, *CST7*, *CCL5* | 1.242 | 0.0073 |
| *LPL* | *APOE*, *TREM2*,*CST7*, *CCL5* | 0.927 | 0.034 |
| *CST7* | *APOE*, *TREM2*, *LPL*, *CCL5* | 1.187 | 0.0067 |
| *CCL5* | *APOE*, *TREM2*, LPL, *CST7* | 1.362 | 0.0041 |

**Panel c | Gene-level *Z* of DAM5 components**

| Gene | *Z* | *P* | NSNPS |
|---|---|---|---|
| *APOE* | 7.53 | $2.6\times10^{-14}$ | 49 |
| *LPL* | 1.95 | 0.026 | 260 |
| *TREM2* | 1.75 | 0.040 | 196 |
| *CST7* | −0.55 | 0.708 | 220 |
| *CCL5* | −0.66 | 0.746 | 66 |

**Panel d | REPAIR12 mapping and attrition status through the MAGMA pipeline (NCBI37.3)**

| Original genes (12) | NCBI37.3 symbol mapping | Genotype panel | Final status |
|---|---|---|---|
| *APOE* | mapped, with SNP count （348） | mapped, with SNP count | Tested |
| *LGALS3BP* | mapped, with SNP count （3959） | mapped, with SNP count | Tested |
| *HLA-A* | not mapped | N/A | not mapped (MHC-I allelic gene naming) |
| *IFITM3* | mapped, with SNP count （10410） | mapped, with SNP count | Tested |
| *HLA-B* | not mapped | N/A | not mapped (MHC-I allelic gene naming) |
| *IFI27* | mapped, with SNP count （3429） | mapped, with SNP count | Tested |
| *RPS21* | mapped, with SNP count （6227） | mapped, with SNP count | Tested |
| *RPL36A* | mapped, with SNP count （6173） | not mapped | not in genotype panel |
| *CCL5* | mapped, with SNP count （6352） | mapped, with SNP count | Tested |
| *CPLX2* | mapped, with SNP count （10814） | mapped, with SNP coun | Tested |
| *GABRA3* | mapped, with SNP count | not mapped | not in genotype panel |

| Original genes (12) | NCBI37.3 symbol mapping | Genotype panel | Final status |
|---|---|---|---|
| | （2556） | | |
| *FCGR2B* | mapped, with SNP count （2213） | mapped, with SNP count | Tested |
| effective genes | 10/12 | 8/10 | 8 |

Note: RPL36A and GABRA3 are absent from the genotype panel and are also non-expressed genes in the GSE121654 subset; their absence has independent causes and is presented here as a parallel fact rather than a causal link. HLA-A and HLA-B cannot be mapped because NCBI37.3 uses allelic naming for MHC-I genes (experimentally verified). REPAIR8 is the effective set entering MAGMA testing; the 8/7/6 effective-gene counts reported derive from that dataset's annotation and expression constraints, consistent with the 8 effective genes in this study.

**Supplementary Table S15 | Four-group coupling summary and sensitivity analyses of the 5xFAD dataset**

**Panel a | Plate-level results of the four groups**

| Group | Plates | Cells | Plate-level mean *r* ± s.e.m. | Positive plates | t statistic | *P* | DAM⁺ proportion |
|---|---|---|---|---|---|---|---|
| 5XFAD | 31 | 3,968 | 0.447±0.031 | 30/31 | t(30)=13.26 | $4.4\times10^{-14}$ | 23.7% |
| C57BL/6（WT） | 28 | 4,869 | 0.195±0.024 | 28/28 | t(27)=7.04 | $1.4\times10^{-7}$ | 1.8% |
| Trem2KO_5XFAD | 6 | not provided per group | 0.318±0.031 | 6/6 | t(5)=9.71 | $2.0\times10^{-4}$ | 2.8% |
| Trem2KO | 3 | not provided per group | 0.084±0.020 | 3/3 | t(2)=4.13 | 0.054 | 0.15% |

Note: Between-group comparisons use plate-level Mann–Whitney tests; 5xFAD versus C57BL/6 gives $P < 10^{-4}$, Cohen's $d = 1.68$. The pure Trem2KO group comprises only 3 plates ($P = 0.054$ under the t(2) convention) and was not formally tested against WT. Complete plate-level r values are provided as an accompanying Supplementary Data table. The s.e.m. column shows logical per-plate pooled-SD-normalized SEM for display, not the unadjusted per-plate statistic; one-sample t-derived plate-level r SEs are 0.0337 (5xFAD) and 0.0277 (C57BL/6), versus the displayed ±0.031 (0.031, 0.024) — an ~8% discrepancy, indicating the displayed ± values were independently rounded and the two are not interchangeable. Between-group comparisons should rely on plate-level Mann–Whitney tests and Cohen's d (pooled plate-level SD as denominator, computed from unrounded per-plate values, d = 1.68). Cell counts for Trem2KO_5xFAD and Trem2KO were not provided per group in the source data; only plate-level statistics are reported.

**Panel b | Composition sensitivity**

| Specification | 5xFAD plate-level *r* | C57BL/6 plate-level *r* | Between-group *P* |
|---|---|---|---|
| all cells | 0.447 | 0.195 | $<10^{-4}$ |
| excluding the top 10% DAM-scoring cells | 0.212 | 0.157 | 0.0032 |

**Panel c | Age sensitivity (6-month matched)**

| Specification | 5xFAD plate-level *r*（positive plates） | C57BL/6 plate-level *r*（positive plates） | Between-group *P* |
|---|---|---|---|
| all ages | 0.447（30/31） | 0.195（28/28） | $<10^{-4}$ |
| 6-month only | 0.475（27/27） | 0.168（19/19） | $<10^{-4}$ |

Note: The 567 cells from 20-month-old C57BL/6 mice do not enter the primary analysis and are noted here only. The dataset metadata lack animal and litter identifiers; age information is partially missing (18 plates).

**Supplementary Table S16 | 95% confidence intervals of coupling change magnitudes (Δ) across systems**

| System | Δ | 95%CI（r scale） | Computational specification |
|---|---|---|---|
| facial nerve axotomy（GSE90975, cell level） | +0.138 | −0.009 - +0.282 | Fisher z difference of the pooled r between the two sides (contra 0.085, ipsi 0.223) |
| LPC demyelination (GSE121654, cell level) | +0.124 | +0.058 - +0.190 | Fisher z difference of the pooled r between the two groups (SALINE 0.296, LPC 0.420) |
| 5xFAD (GSE98969, plate level) | +0.252 | +0.166 - +0.338 | difference of plate-level means; SE composed from implied SEMs of two one-sample t statistics |
| Trem2KO_5XFAD (plate level) | +0.123 | +0.039 - +0.207 | same as above, control = C57BL/6 (0.195) |
| SEA-AD (donor level) | +0.072 | spans zero (see note) | Fisher z difference of Braak-binned within-r means (Early n=6, Late n=49) |

**Note:** Δ and the 95%CIs are on the r scale; SEs were composited from z-scale differences and back-transformed for approximation. The SEs for the 5xFAD and Trem2KO_5xFAD rows derive from the implied SEMs of one-sample t statistics in Supplementary Table 15 (0.0337, 0.0328, and 0.0277). For SEA-AD, the Early group comprises only 6 donors and the Fisher *z* CI is approximately −0.80 to +0.85, uninformative; hence the 95%CI is labeled “spans zero”, with binned testing (Early vs Late, Mann–Whitney $P = 0.13$) and weighted regression ($b = 0.0112$, $P = 0.054$) provided as references; the Δ in this row serves cross-magnitude comparison only. The interval spanning zero for the facial nerve axotomy row is consistent with its exploratory designation.

**Supplementary Table S17 | Evidence grading of each component of the working model**

| Component | Evidence level | Primary evidence | Method and limitations |
|---|---|---|---|
| phagocytic load (injury background) | 0 | injury models; injury-side phagocytosis and lipid metabolism module elevation | no direct measurement, background premise |
| lysosomal mechanical stress | 0 | — | no functional readout of membrane tension or mechanosensitive channels |
| TMEM63A / TRPML1 | 1 | TMEM63A 22.1% and MCOLN1 12.2% transcripts detectable | no functional readout of membrane tension or mechanosensitive channels |
| MiT/TFE transcriptional response | 3 | lysosomal program peaks at day 7 (1.350), earlier than the DAM differential peak (day 30) | population-mean dynamics, not single-cell trajectory inference |
| TREM2 dependence | 3 | 5xFAD Δ=+0.252; Trem2KO_5xFAD Δ=+0.318; DAM$^{+}$ proportion 23.7% → 2.8% | plate-level statistical units; pure Trem2KO group only 3 plates |
| lysosomal program → DAM coupling | 5 | mouse 0.137 and 0.361; human cell-level 0.096 and 0.203, donor-level 0.534 and 0.491; SEA-AD 0.554; cluster-robust SE and Bonferroni correction | correlational evidence, causal direction undetermined |
| Apoe$^{+}$ repair state | 4 | C1 Apoe$^{+}$ 57.7%；Cohen's d=1.85-1.93；MAGMA APOE | subpopulation definition depends on module scoring |
| genetic anchoring (APOE)） | 5b | MAGMA DAM5 β=1.947，P=8.7×10$^{-6}$；LYS10 P=0.357 | summary statistics exclude 23andMe |

Note: Levels are determined by the available data structure, independent of effect size; level 0 indicates that the component has no direct measurement in this study and serves only as a background premise of the working model.

**Supplementary Table S18 | Sample structure and statistical units of the two murine datasets**

| Dataset | Batch or sample | Time or group | Laterality | Cells |
|---|---|---|---|---|
| GSE90975 (4 sequencing plates, 10 animals, 1,437 cells in total) | P1 | Day 0 | baseline (unlateralized) | 82 |
| | P1 | Day 30 | contralateral | 92 |
| | P1 | Day 7 | contralateral | 96 |
| | P1 | Day 7 | ipsilateral | 96 |
| | P2 | Day 0 | baseline (unlateralized) | 93 |

| Dataset | Batch or sample | Time or group | Laterality | Cells |
|---|---|---|---|---|
| | P2 | Day 30 | contralateral | 79 |
| | P2 | Day 30 | ipsilateral | 173 |
| | P3 | Day 0 | baseline (unlateralized) | 177 |
| | P3 | Day 7 | contralateral | 94 |
| | P3 | Day 7 | ipsilateral | 88 |
| | P4 | Day 30 | contralateral | 94 |
| | P4 | Day 30 | ipsilateral | 93 |
| | P4 | Day 7 | contralateral | 94 |
| | P4 | Day 7 | ipsilateral | 86 |
| GSE121654(6 samples corresponding to 6 independent animals, 5,325 cells in total) | GSM3442041 | SALINE | not applicable | 1159 |
| | GSM3442042 | SALINE | not applicable | 1017 |
| | GSM3442043 | SALINE | not applicable | 410 |
| | GSM3442044 | LPC | not applicable | 1136 |
| | GSM3442045 | LPC | not applicable | 1128 |
| | GSM3442046 | LPC | not applicable | 475 |

Note: In GSE90975, day 0 is the pre-surgical baseline and is not lateralized. In GSE121654, each sample corresponds to one independent animal; all are P100 female mice harvested 7 days post-treatment.

**Supplementary Table S19 | Sample structure and statistical units of the two human cohorts**

| Item | Lee 2026[14] | Rexach 2024[15] |
|---|---|---|
| total donors (analyzed) | 578 (AD 402, control 176) | 38 (AD 10, control 9, PSP 11, Pick’s disease 8) |
| nuclei | 100,379 | 21,572 |
| subcohort or region | MSSM 481、RADC 97 | Insula 16,976, BA4 3,511, V1 1,088 (nuclei) |
| donor × region units | N/A | 82 |
| covariates | Age, sex (interval-censored) | Age (60–79-year bracket, 61%), sex, brain region |
| effective genes for scoring | lys 10/10、dam 5/5、dam4 4/4、rep 12/12 | GBA corresponds to GBA1 in this dataset’s annotation |
| ≥20 nuclei per donor threshold | Yes | Yes |

Note: In the Rexach cohort, nuclei before the ≥20-nuclei-per-donor threshold were 21,575 (insula 16,976; BA4 3,511; V1 1,088); after thresholding, 38 donors and 21,572 nuclei were analyzed. The in-table nuclei and region counts correspond to the pre- and post-threshold conventions, respectively (Sections 2.3.8 and Supplementary Table 14).

**Supplementary Table S20 | SEA-AD cohort summary**

**Panel a | Donor-level summary (n = 84)**

| Donor ID | Braak | Age | Sex | Nuclei | Lysosomal | DAM | Depth |
|---|---|---|---|---|---|---|---|
| H19.33.004 | 4 | 80 | Female | 2874 | 0.119 | -0.027 | 6136 |
| H20.33.001 | 4 | 82 | Male | 4644 | 0.132 | 0.04 | 7246 |
| H20.33.002 | 4 | 97 | Female | 1789 | 0.091 | 0.004 | 7416 |
| H20.33.004 | 5 | 86 | Male | 1409 | 0.31 | 0.102 | 7025 |
| H20.33.005 | 4 | 99 | Female | 3064 | 0.072 | -0.093 | 5908 |
| H20.33.008 | 5 | 92 | Female | 5257 | 0.029 | -0.073 | 7631 |
| H20.33.011 | 5 | 93 | Female | 3897 | 0.079 | 0.007 | 6495 |
| H20.33.012 | 3 | 91 | Female | 3783 | 0.13 | -0.049 | 7535 |
| H20.33.013 | 4 | 94 | Male | 3249 | 0.107 | -0.052 | 5290 |
| H20.33.014 | 4 | 82 | Female | 1254 | 0.043 | -0.108 | 5585 |
| H20.33.015 | 5 | 88 | Male | 5489 | 0.194 | 0.051 | 6979 |
| H20.33.016 | 4 | 93 | Female | 4408 | 0.079 | -0.031 | 5064 |
| H20.33.017 | 6 | 69 | Male | 5068 | 0.063 | 0.076 | 7010 |
| H20.33.018 | 6 | 81 | Female | 3407 | 0.176 | 0.096 | 6080 |
| H20.33.019 | 3 | 87 | Female | 3725 | 0.091 | -0.061 | 7042 |
| H20.33.020 | 6 | 81 | Male | 6400 | 0.143 | -0.027 | 6483 |
| H20.33.024 | 5 | 90 | Male | 662 | 0.136 | -0.03 | 4304 |
| H20.33.025 | 5 | 94 | Male | 6885 | 0.091 | 0.027 | 5272 |
| H20.33.026 | 6 | 75 | Female | 5091 | 0.117 | 0.029 | 6034 |
| H20.33.027 | 4 | 99 | Female | 2823 | 0.144 | 0.001 | 4461 |
| H20.33.028 | 5 | 94 | Female | 5601 | 0.179 | -0.015 | 5042 |
| H20.33.029 | 5 | 91 | Female | 4740 | 0.093 | -0.06 | 6396 |
| H20.33.030 | 5 | 86 | Female | 1218 | 0.127 | -0.031 | 6843 |
| H20.33.031 | 6 | 87 | Female | 4470 | 0.09 | 0.012 | 6864 |
| H20.33.032 | 5 | 98 | Male | 2277 | 0.099 | -0.019 | 5101 |
| H20.33.033 | 6 | 68 | Male | 4002 | 0.119 | -0.036 | 6952 |
| H20.33.034 | 4 | 85 | Female | 687 | 0.214 | 0.108 | 4564 |
| H20.33.035 | 4 | 99 | Female | 3736 | 0.152 | 0.043 | 6162 |
| H20.33.036 | 5 | 100 | Female | 2257 | 0.191 | -0.014 | 6256 |
| H20.33.037 | 5 | 96 | Female | 1088 | 0.216 | -0.054 | 7917 |
| H20.33.038 | 5 | 90 | Female | 966 | 0.187 | 0.029 | 6165 |
| H20.33.039 | 5 | 96 | Female | 3817 | 0.086 | -0.02 | 5039 |
| H20.33.040 | 4 | 98 | Male | 5860 | 0.128 | 0.132 | 5551 |
| H20.33.041 | 5 | 91 | Female | 837 | 0.234 | 0.045 | 5197 |
| H20.33.043 | 4 | 85 | Male | 1694 | 0.286 | 0.148 | 5657 |
| H20.33.044 | 5 | 81 | Male | 1086 | 0.218 | -0.041 | 4200 |
| H20.33.045 | 6 | 77 | Female | 1121 | 0.172 | 0.104 | 5532 |
| H20.33.046 | 6 | 94 | Male | 3536 | 0.045 | -0.072 | 4859 |
| H21.33.001 | 3 | 80 | Male | 4977 | 0.165 | 0.116 | 4787 |
| H21.33.002 | 6 | 70 | Female | 4761 | 0.087 | 0.141 | 7710 |
| H21.33.003 | 0 | 78 | Male | 3772 | 0.022 | -0.129 | 5606 |
| H21.33.004 | 2 | 93 | Male | 3834 | 0.071 | -0.048 | 5150 |
| H21.33.005 | 5 | 95 | Male | 1235 | 0.073 | -0.095 | 4926 |

| Donor ID | Braak | Age | Sex | Nuclei | Lysosomal | DAM | Depth |
|---|---|---|---|---|---|---|---|
| H21.33.006 | 4 | 97 | Male | 1441 | 0.089 | -0.031 | 5720 |
| H21.33.007 | 5 | 86 | Female | 1303 | 0.102 | -0.026 | 6789 |
| H21.33.008 | 5 | 91 | Female | 1705 | 0.169 | -0.055 | 7093 |
| H21.33.009 | 6 | 65 | Female | 487 | 0.224 | -0.017 | 8605 |
| H21.33.010 | 6 | 93 | Female | 1469 | 0.24 | 0.134 | 7058 |
| H21.33.011 | 4 | 83 | Female | 2108 | 0.106 | -0.072 | 6573 |
| H21.33.012 | 4 | 93 | Female | 1330 | 0.198 | 0.156 | 5308 |
| H21.33.013 | 5 | 94 | Female | 2193 | 0.154 | 0.086 | 7108 |
| H21.33.014 | 4 | 92 | Male | 1251 | 0.243 | 0.173 | 6296 |
| H21.33.015 | 4 | 98 | Male | 2771 | 0.172 | -0.001 | 6839 |
| H21.33.016 | 4 | 94 | Female | 1459 | 0.168 | 0.075 | 5357 |
| H21.33.017 | 5 | 92 | Female | 2372 | 0.134 | -0.106 | 9053 |
| H21.33.018 | 2 | 89 | Female | 4790 | 0.146 | 0.031 | 5792 |
| H21.33.019 | 0 | 75 | Male | 2816 | 0.135 | -0.079 | 4616 |
| H21.33.020 | 5 | 82 | Male | 445 | 0.415 | 0.15 | 3527 |
| H21.33.021 | 3 | 99 | Male | 591 | 0.177 | 0.047 | 7346 |
| H21.33.022 | 6 | 82 | Female | 4071 | 0.132 | -0.006 | 6950 |
| H21.33.023 | 4 | 102 | Male | 2549 | 0.076 | -0.003 | 8013 |
| H21.33.025 | 4 | 88 | Female | 2569 | 0.142 | -0.074 | 6770 |
| H21.33.026 | 5 | 90 | Female | 4281 | 0.111 | -0.018 | 7658 |
| H21.33.027 | 5 | 92 | Male | 5580 | 0.109 | 0.119 | 4731 |
| H21.33.028 | 2 | 72 | Male | 7205 | 0.09 | -0.043 | 9127 |
| H21.33.029 | 5 | 89 | Male | 2300 | 0.124 | 0.104 | 5957 |
| H21.33.030 | 3 | 89 | Male | 1308 | 0.15 | 0.03 | 5972 |
| H21.33.031 | 5 | 84 | Male | 1860 | 0.147 | 0.026 | 5493 |
| H21.33.032 | 4 | 98 | Female | 4508 | 0.118 | 0.038 | 5064 |
| H21.33.033 | 5 | 83 | Female | 3476 | 0.09 | -0.058 | 6160 |
| H21.33.034 | 6 | 90 | Female | 625 | 0.21 | 0.066 | 6064 |
| H21.33.035 | 5 | 97 | Female | 4195 | 0.144 | 0.047 | 7899 |
| H21.33.036 | 5 | 93 | Female | 3052 | 0.086 | 0.035 | 5712 |
| H21.33.037 | 4 | 88 | Female | 1268 | 0.076 | -0.013 | 7391 |
| H21.33.038 | 3 | 84 | Female | 3156 | 0.114 | -0.058 | 6124 |
| H21.33.039 | 5 | 88 | Female | 1872 | 0.12 | -0.046 | 6622 |
| H21.33.040 | 5 | 83 | Male | 954 | 0.197 | 0.087 | 5696 |
| H21.33.041 | 4 | 98 | Female | 1044 | 0.16 | 0.024 | 5198 |
| H21.33.042 | 5 | 91 | Female | 1619 | 0.139 | -0.051 | 7218 |
| H21.33.043 | 2 | 95 | Female | 1497 | 0.149 | 0.046 | 5435 |
| H21.33.044 | 6 | 88 | Female | 1629 | 0.301 | 0.099 | 6730 |
| H21.33.045 | 6 | 94 | Female | 1349 | 0.225 | 0.123 | 6384 |
| H21.33.046 | 5 | 97 | Male | 1665 | 0.154 | 0.135 | 7818 |
| H21.33.047 | 5 | 90 | Male | 1089 | 0.116 | -0.026 | 6945 |

**Panel b | Per-donor within-r and Braak stratification summary**

| Donor ID | Braak | Within-*r* | Within-*r* (*z*-mean) | Compressio n | Nuclei |
|---|---|---|---|---|---|
| H19.33.004 | 4 | 0.249 | | | 2874 |
| H20.33.001 | 4 | 0.297 | | | 4644 |
| H20.33.002 | 4 | 0.233 | | | 1789 |

| **Donor ID** | **Braak** | **Within-*r*** | **Within-*r* (*z*-mean)** | **Compressio n** | **Nuclei** |
|---|---|---|---|---|---|
| H20.33.004 | 5 | 0.428 | | | 1409 |
| H20.33.005 | 4 | 0.29 | | | 3064 |
| H20.33.008 | 5 | 0.261 | | | 5257 |
| H20.33.011 | 5 | 0.25 | | | 3897 |
| H20.33.012 | 3 | 0.356 | | | 3783 |
| H20.33.013 | 4 | 0.278 | | | 3249 |
| H20.33.014 | 4 | 0.206 | | | 1254 |
| H20.33.015 | 5 | 0.416 | | | 5489 |
| H20.33.016 | 4 | 0.281 | | | 4408 |
| H20.33.017 | 6 | 0.34 | | | 5068 |
| H20.33.018 | 6 | 0.36 | | | 3407 |
| H20.33.019 | 3 | 0.363 | | | 3725 |
| H20.33.020 | 6 | 0.274 | | | 6400 |
| H20.33.024 | 5 | 0.027 | | | 662 |
| H20.33.025 | 5 | 0.248 | | | 6885 |
| H20.33.026 | 6 | 0.373 | | | 5091 |
| H20.33.027 | 4 | 0.208 | | | 2823 |
| H20.33.028 | 5 | 0.336 | | | 5601 |
| H20.33.029 | 5 | 0.322 | | | 4740 |
| H20.33.030 | 5 | 0.326 | | | 1218 |
| H20.33.031 | 6 | 0.302 | | | 4470 |
| H20.33.032 | 5 | 0.206 | | | 2277 |
| H20.33.033 | 6 | 0.325 | | | 4002 |
| H20.33.034 | 4 | 0.224 | | | 687 |
| H20.33.035 | 4 | 0.207 | | | 3736 |
| H20.33.036 | 5 | 0.386 | | | 2257 |
| H20.33.037 | 5 | 0.267 | | | 1088 |
| H20.33.038 | 5 | 0.342 | | | 966 |
| H20.33.039 | 5 | 0.208 | | | 3817 |
| H20.33.040 | 4 | 0.322 | | | 5860 |
| H20.33.041 | 5 | 0.306 | | | 837 |
| H20.33.043 | 4 | 0.204 | | | 1694 |
| H20.33.044 | 5 | 0.272 | | | 1086 |
| H20.33.045 | 6 | 0.314 | | | 1121 |
| H20.33.046 | 6 | 0.29 | | | 3536 |
| H21.33.001 | 3 | 0.16 | | | 4977 |
| H21.33.002 | 6 | 0.234 | | | 4761 |
| H21.33.003 | 0 | 0.218 | | | 3772 |
| H21.33.004 | 2 | 0.275 | | | 3834 |
| H21.33.005 | 5 | 0.253 | | | 1235 |
| H21.33.006 | 4 | 0.331 | | | 1441 |
| H21.33.007 | 5 | 0.275 | | | 1303 |
| H21.33.008 | 5 | 0.411 | | | 1705 |
| H21.33.009 | 6 | 0.312 | | | 487 |
| H21.33.010 | 6 | 0.433 | | | 1469 |
| H21.33.011 | 4 | 0.165 | | | 2108 |
| H21.33.012 | 4 | 0.274 | | | 1330 |
| H21.33.013 | 5 | 0.357 | | | 2193 |
| H21.33.014 | 4 | 0.34 | | | 1251 |

| Donor ID | Braak | Within-*r* | Within-*r* (*z*-mean) | Compression | Nuclei |
|---|---|---|---|---|---|
| H21.33.015 | 4 | 0.215 | | | 2771 |
| H21.33.016 | 4 | 0.327 | | | 1459 |
| H21.33.017 | 5 | 0.256 | | | 2372 |
| H21.33.018 | 2 | 0.273 | | | 4790 |
| H21.33.019 | 0 | 0.319 | | | 2816 |
| H21.33.020 | 5 | 0.299 | | | 445 |
| H21.33.021 | 3 | 0.385 | | | 591 |
| H21.33.022 | 6 | 0.291 | | | 4071 |
| H21.33.023 | 4 | 0.358 | 0.231 | 0.139 | 2549 |
| H21.33.025 | 4 | 0.269 | | | 2569 |
| H21.33.026 | 5 | 0.288 | | | 4281 |
| H21.33.027 | 5 | 0.217 | | | 5580 |
| H21.33.028 | 2 | 0.31 | | | 7205 |
| H21.33.029 | 5 | 0.32 | | | 2300 |
| H21.33.030 | 3 | 0.318 | | | 1308 |
| H21.33.031 | 5 | 0.257 | | | 1860 |
| H21.33.032 | 4 | 0.21 | | | 4508 |
| H21.33.033 | 5 | 0.258 | | | 3476 |
| H21.33.034 | 6 | 0.362 | | | 625 |
| H21.33.035 | 5 | 0.33 | | | 4195 |
| H21.33.036 | 5 | 0.261 | | | 3052 |
| H21.33.037 | 4 | 0.252 | | | 1268 |
| H21.33.038 | 3 | 0.302 | | | 3156 |
| H21.33.039 | 5 | 0.34 | | | 1872 |
| H21.33.040 | 5 | 0.412 | | | 954 |
| H21.33.041 | 4 | 0.267 | | | 1044 |
| H21.33.042 | 5 | 0.355 | | | 1619 |
| H21.33.043 | 2 | 0.188 | 0.129 | 0.069 | 1497 |
| H21.33.044 | 6 | 0.439 | 0.324 | 0.116 | 1629 |
| H21.33.045 | 6 | 0.385 | | | 1349 |
| H21.33.046 | 5 | 0.416 | | | 1665 |
| H21.33.047 | 5 | 0.208 | | | 1089 |

**Bin-level summary (coarse Braak bins)**

| Braak bin | Donors | Mean | SD | Median |
|---|---|---|---|---|
| Early (Braak 0-II) | 6 | 0.264 | 0.051 | 0.274 |
| Mid (Braak III-IV) | 29 | 0.272 | 0.062 | 0.274 |
| Late (Braak V-VI) | 49 | 0.309 | 0.076 | 0.312 |

**Panel c | Sensitivity panel**

| Analysis | Result | Definition |
|---|---|---|
| Main donor-level r | r = 0.554 (95% CI 0.385-0.686), *P* = 4.7e-8, *n* = 84 | Pearson r across donor means (lysosomal vs DAM) |
| Depth adjustment | partial r = 0.544, retained 98.3% | Residualised on log nuclei per donor |
| WLS Braak (main) | b = 0.0112, *P* = 0.054 | within-z on Braak, age, sex, log |

| Analysis | Result | Definition |
|---|---|---|
| | | nuclei; weighted by nuclei |
| WLS, excluding Braak 0 | b = 0.0123, *P* = 0.082 | 2 donors removed |
| WLS, excluding Severely Affected | b = 0.0107, *P* = 0.095 | 11 donors removed |
| Braak squared term | b = 0.0040, *P* = 0.250 | No non-linear evidence |
| Composition | DAM-high fraction vs Braak: *r* = 0.178, *P* = 0.105 | DAM top-10% cell fraction per donor |
| Bins, Early vs Mid | Mann-Whitney P = 0.78 | *n* = 6 vs 29 |
| Bins, Early vs Late | Mann-Whitney P = 0.13 | *n* = 6 vs 49 |
| Bins, Mid vs Late | Mann-Whitney P = 0.021 (Bonferroni x3 = 0.062) | *n* = 29 vs 49 |
| Implementation, cell level | chunked vs scanpy r = 0.984 (lysosomal), 0.990 (DAM) | Scoring implementation check |
| Implementation, within-r | z-mean compression 0.069-0.139 | Per-donor scanpy minus z-mean within-r |
| Braak vs lysosomal | *r* = 0.201, *P* = 0.067 | Descriptor, no significant trend |
| Braak vs DAM | *r* = 0.234, *P* = 0.032 | Descriptor |
| Braak vs within-r | *r* = 0.238, *P* = 0.030 | Nominal only, not adjusted |

Note: within-r denotes the Pearson correlation between the lysosomal program and DAM scores within each donor; *z*-mean denotes the same-quantity mean after per-donor standardization; "Compression" is their difference (0.07–0.14). Panel C P-values are two-sided and uncorrected unless otherwise noted; binned tests are Mann–Whitney U tests with Bonferroni correction across three comparisons.

**Supplementary Table S21 | Consistency test of donor-inclusion thresholds in the Rexach cohort**

| Metric | ≥20 nuclei (38 donors, 21,572 nuclei) | ≥50 nuclei (35 donors, 21,452 nuclei) | Consistency assessment |
|---|---|---|---|
| cell-level pooled r (95%CI) | +0.174 (+0.161 - +0.187) | +0.174 (+0.161 - +0.187) | consistent (\|Δr\| = 0.0004) |
| pooled P | $2.9\times10^{-146}$ | $3.7\times10^{-146}$ | consistent |
| donor-mean *r* | +0.379 | +0.510 | direction and significance concordant, magnitude shift +0.131 |
| donor-level P | 0.019 | 0.002 | consistent |
| included nuclei | 21,572(pre-threshold 21,575) | 21,452 (99.4% of the ≥20-nuclei subset) | — |

Note: ≥20 nuclei is the primary threshold; ≥50 nuclei is the sensitivity threshold. Cell-level pooled correlations are donor-stratified Fisher z-weighted; donor-mean correlations are Pearson correlations of 38 and 35 donor means respectively. Cell-level estimates agree exactly under both thresholds; donor-level direction and significance are concordant, so the primary result is robust to threshold choice. The +0.131 donor-mean drift originates from the small n = 35 sample and removal of 3 small donors, with the direction of enhancement; this indicates that small donors introduce measurement noise rather than genuine heterogeneity.
